# TriTan: An efficient triple non-negative matrix factorisation method for integrative analysis of single-cell multiomics data

**DOI:** 10.1101/2023.07.14.549059

**Authors:** Xin Ma, Lijing Lin, Qian Zhao, Mudassar Iqbal

**Affiliations:** Division of Informatics, Imaging and Data Sciences, Faculty of Biology, Medicine and Health, University of Manchester, Manchester, U.K.; Translation Manchester, Faculty of Biology, Medicine and Health, University of Manchester, Manchester, U.K.

## Abstract

**Motivation:** Single-cell multi-omics have opened up tremendous opportunities for understanding gene regulatory networks underlying cell states by simultaneously profiling transcriptomes, epigenomes and proteomes of the same cell. However, existing computational methods for integrative analysis of these high-dimensional multi-modal data are either computationally expensive or limited in interpretation ans scope. These limitations pose challenges in the implementation of these methods in large-scale studies and hinder a more in-depth understanding of the underlying regulatory mechanisms.

**Results:** Here, we propose TriTan (Triple inTegrative fast non-negative matrix factorisation), an efficient joint factorisation method for single-cell multiomics data. TriTan implements a highly efficient triple non-negative matrix factorisation algorithm which greatly enhances its computational speed, and facilitates interpretation by clustering both the cells and features simultaneously as well as identifying signature feature sets for each cell cluster. Additionally, three matrix factorisation produced by TriTan helps in finding associations of features across modalities, facilitating the prediction of cell type specific regulatory networks. We applied TriTan to single-cell multi-modal data obtained from different technologies and benchmarked it against the state-of-the-art methods where it shows highly competitive performance. Furthermore, we showed a range of downstream analyses that can be conducted utilising the outputs from TriTan.

**Availability:** https://github.com/maxxxxxxxin/TriTan online.

## Introduction

Single-cell multi-modal technologies can simultaneously profile multiple omics modalities within the same cell, including transcriptomics, epigenomics, and proteomics (1). These approaches provide a more comprehensive view of cellular function as they capture multiple layers of regulatory machinery of the cell. Integration of epigenomics and proteomics data with transcriptomics information can reveal new insights into the underlying mechanisms that regulate gene expression and how these may alter in response to different stimuli.

With the rapid advancement in these technologies, we are faced with a complex data integration challenge. The unique characteristics of data from each modality, such as sparsity, dimension, and scale, make it difficult to synthesize a coherent joint representation. In order to effectively combine information from different modalities and generate meaningful and interpretable signatures, it is necessary to design bespoke methodologies that are capable of seamlessly integrating multi-modal data and are flexible to include more modalities as they become available.

Several recent studies have offered innovative ways of integrating single-cell multiomics data. One category of methods focus on finding relationships among data from different modalities and then perform the integration. For example, Seurat v4 (2) uses Weighted Nearest Neighbors (WNN) approach to learn a cell-specific weight and then integrate all modalities. Schema (3) uses a metric learning strategy to extract informative features for each modality and integrate them into a single coherent interpretation. Another approach consists of searching for a common low-dimensional feature latent space and then projecting different views into the latent spaces where subsequent clustering tasks can be conducted. In this category, MOFA+ (4) jointly models variations across multiple modalities after low dimensional reconstructions using a factor modelling approach with variational inference. scAI (5) method infers the low-dimensional representations via a matrix factorisation model and UINMF (6) derives a non-negative matrix factorisation algorithm for integrating modalities with an additional matrix that allows un-shared features to inform the factorisation. Recently published, MOJITOO (7) efficiently detects multiple modalities’ shared representations through canonical correlation analysis (CCA). Besides, deep generative models have been employed to perform single-cell multiomics data integration. Among these, totalVI (8) uses a variational autoencoder (VAE) model to jointly analyse CITE-seq data (RNA and cell surface proteins from same cell). It represents the data as a composite of biological and technical factors, including protein background and batch effects. Of note, totalVI only allows integration of particular modalities (scRNA-seq and protein abundance). CLUE (9) uses multi-modal VAEs and considers cross-encoders to construct latent representations from modality-incomplete observations.

The majority of the methods discussed above have a primary focus on cell type identification. Among them, matrix factorisation methods have become popular as they provide low dimensional representation of the data as well as interpretable feature contributions for the discovered topics. However, standard two-matrix factorisation methods suffer from several limitations, including arbitrariness in determining the number of cell topics and importantly no systematic way to find signature features for each cell topic and associations of features across different modalities. Below we describe our non-negative matrix factorisation method, TriTan, which aims to address above limitations.

## Approach

In the context of single-cell multiomics data, the critical need to find the functional roles of features in individual modalities and the associations between them motivates us to look beyond the existing two-matrix factorisation methods. We have designed TriTan, an efficient and interpretable method that decomposes the input single-cell multi-modal matrices into following low-dimensional matrices: a shared cell cluster matrix across all modalities, distinct feature-cluster matrices, and association matrices that capture the associations between cell clusters and feature clusters for each modality (see cartoon in Fig. 1). TriTan enables the simultaneous detection of latent cell clusters and feature clusters, as well as the exploration of associations between features, such as the links between genes and potential regulatory peaks.

**Fig. 1.**
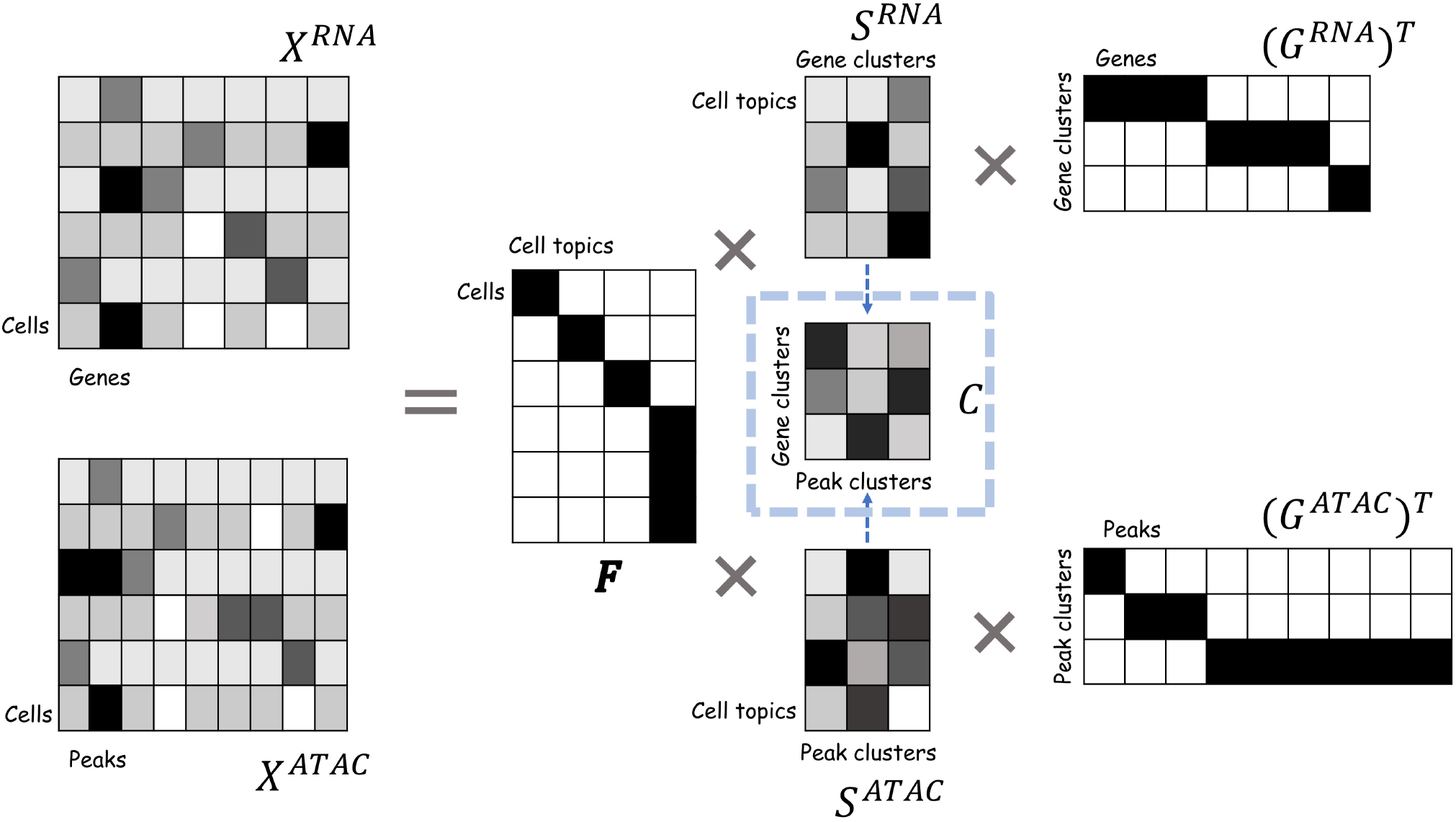
Schematic of TriTan: It receives two (or more) paired single-cell matrices whose rows represent the same cells, where each matrix represents a particular molecular modality. In this example, scRNA and scATAC-seq, are being considered. Each input matrix is decomposed into three matrices where *F* is a cell cluster matrix shared across all modalities. Each column of *F* represents a distinct cell cluster and each row contains a single non-zero element indicating the cluster assignment of the corresponding cell. *G*^RNA^ and *G*^ATAC^ are feature cluster matrices whose columns represent different feature clusters and rows contain a single non-zero element indicating the cluster assignment of the corresponding feature. Middle matrices *S*^RNA^ and *S*^ATAC^ (association matrices) are condensed representations of the input matrices by cell clusters and feature clusters, which can be considered as associations between cell clusters and feature clusters. After the factorisation, we calculated the matrix *C* using *S*^RNA^ and *S*^ATAC^ to show feature correlations across modalities.

Unlike traditional matrix factorisation methods, TriTan does not rely on predefined parameters, such as the rank of factor matrices. TriTan uses an automated approach for identifying the number of cell and feature clusters, making it a more flexible and adaptive tool for analysing complex datasets. TriTan is implemented in Python and takes multi-modal data format (MuData) (10) as input and it is compatible with popular platforms such as Scanpy(11).

We benchmarked TriTan against other state-of-the-art methods, Seurat v4 (WNN), MOFA+ and MOJITOO, using three public multi-modal datasets with two modalities (scRNA-seq and scATAC-seq) in terms of their ability to recover known cell clusters as well as computational efficiency and memory requirements. Moreover, through the association matrices, TriTan offers a unique perspective on the relationships between different types of omics data within individual cells and facilities downstream studies of underlying regulatory mechanisms.

## Materials and methods

### A. TriTan

TriTan is designed to deal with the following problem: A multiomics dataset 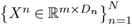 includes *N* single-cell data from different modalities, where *m* represents the number of cells shared across modalities, and *D*_*n*_ represents the number of features in modality *n*. We aim to discover *K*_1_ cell clusters that are shared across all modalities, and simultaneously, find 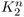 feature clusters for the *n*-th modality, *n* = 1, …, *N*. Note that the number of cell topics, *K*_1_, does not need to be predetermined but rather learned by TriTan. Additionally, TriTan also finds modality specific association matrices which reflects the weights of each feature cluster across all cell clusters. In summary, TriTan learns a representation for each modality : *X*^*n*^ ≈ *FS*^*n*^(*G*^*n*^)^*T*^, where *F* is a binary matrix, shared across all modalities, with a single non-zero element on each row indicating the cluster assignment of the corresponding cell; *G*^*n*^ is a binary feature cluster matrix with each row containing a single non-zero element indicating the cluster assignment of the corresponding feature; *S*^*n*^ is the modality specific association matrix. In the following, ‖*·*‖ refers to the norm ‖*A*‖^2^ = Tr(*AA*^*T*^), and *A*_*i*·_ and *A*_·*j*_ denote the *i*-th row and *j*-th of column of *A*, respectively.

#### A.1. An efficient multiplicative update algorithm for Non-negative Matrix Tri-factorisation (NMTF)

Traditional NMTF methods (12) which infer three non-negative latent matrices based on Multiplicative Update (MU) (13) suffer from slow computational speed and considerable memory consumption. To accelerate NMTF on large-scale data, we design a novel optimization algorithm with motivation to speed up the factorisation. Firstly, to set the scene, we present our factorisation scheme for the case of a single matrix *X*. Here, the aim is to minimise the following loss function:

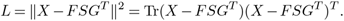

*L* can be minimised over *F, G* and *S* in an alternating fashion. Since F and G are aimed to produce the cluster assignments for cells and features, respectively, the computation of *F* and *G* can be simplified. With fixed *S* and *G*, letting *U* = *SG*^*T*^, in order to minimise 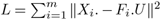, it suffices to minimise ‖*X*_*i*_. − *F*_*i*.*U*_ ‖^2^ for each row *i* of *X*. Since *F*_*i*_. has only one non-zero element, this is equivalent to min_*k*_ ‖*X*_*i*_. − *U*_*k*_. ‖^2^. Therefore, for each row *i* of *F* :

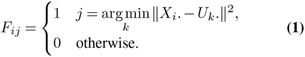

Computation of *G* with fixed *F* and *S* uses the same strategy as the above. With *V* = *FS*, for each column *j* of *X*, noted as *X*_.*j*_, we aim to minimise ‖*X*_.*j*_ − *V* (*G*^*T*^)_.*j*_ ‖^2^. Since (*G*^*T*^)_.*j*_ = *G*_*j*._ and it only has one non-zero element, this is equivalent to min_*k*_ ‖*X*_.*j*_ − *V*_.*k*_ ‖^2^. Therefore, for each row *i* of *G*:

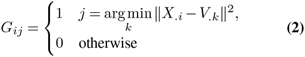

The computation of *S*, given fixed *F* and *G*, uses the following procedure:

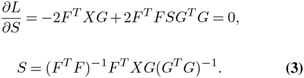

#### A.2. Reducing the dimensions during multiplicative updates

To further improve the efficiency of the factorisation method described above, we propose to use singular value decomposition (SVD) to reduce the dimensions during multiplicative updates. This is mainly designed to speed up the factorisation without losing accuracy even if a small number of dimensions are used. Besides, it helps to reduce the imbalance in the number of features between different modalities (for instance, the number of features can vary significantly between scRNA-seq data and scATAC-seq data).

For a given modality *n*, input matrix *X*^*n*^ (of dimension *m × D*_*n*_) is firstly factorised into three matrices using SVD: *X*^*n*^ = *P*^*n*^Σ^*n*^(*Q*^*n*^)^*T*^, where (*P*^*n*^)^*T*^ *P*^*n*^ = *I*, (*Q*^*n*^)^*T*^ *Q*^*n*^ = *I* and Σ^*n*^is a rectangular diagonal matrix with non-negative numbers (singular values) on the diagonal. We approximate *X*^*n*^ by a truncated SVD, i.e., 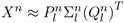 using the first *l* SVD components where *l* can be chosen using a scree plot of the singular values in Σ^*n*^ (14). We use the same *l* for all modalities, and we used different number of components for different datasets using singular values scree plots (see details in supplementary Table. S4). Also in our software package, we provided an option for the user to choose the number of the components for each modality. To reduce computational time and memory usage, *F*^*n*^ and *G*^*n*^ in the previous section are updated with *X*^*n*^ projected to the lower dimension space 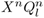 and 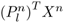, respectively (as shown below in the equations Eq. (4) and Eq. (7)).

#### A.3. Joint multiplicative update for single-cell multiomics data

In previous sections, we described our efficient implementation for NMTF for a single modality. Building on that, here we present TriTan’s joint multiple updates scheme incorporating SVD, to jointly factorise single-cell multi-modal data.

Since there can be significant differences in the data attributes between different modalities, we introduce modality-specific cell weights, 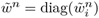, a diagonal matrix with entry 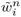 representing the weight for cell *i* in modality *n*. Ideally, we want to optimise the weighted reconstruction loss for all modalities formulated as

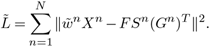

Instead of assigning weights to each modality based on prior knowledge, we adopt an alternative strategy, learning the modality-specific cell weights within the joint update. We introduce an intermediate weighted cell cluster matrix 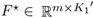 (with a chosen initial number of clusters noted as resolution, *K*_1_′ < *K*_1_), and approximate the weights 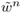 with weights *w*^*n*^ that are updated as a part for updating *F*^⋆^, as shown below in the equations Eq. (4), Eq. (5), and Eq. (6). By doing this, we also learn a comprehensive shared cell representation.

At the (*t* + 1)-th iteration, we firstly update the cell cluster assignment for each modality with ^(*t*)^*U*^*n*^ = ^(*t*)^*S*^*n*^(^(*t*)^*G*^*n*^)^*T*^ :

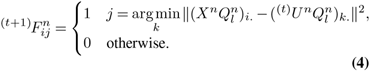

To design a cell-specific weight for each modality, we consider the reconstruction loss for each modality as well as biases that may exist between modalities in a balanced manner (for example, scRNA-seq data usually have much higher average expressions than scATAC-seq data). Here, *w*_*n*_ are updated as follows:

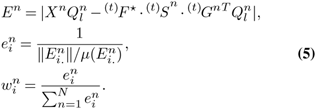

With these weights, a weighted cell cluster assignment is constructed

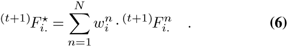

With the computed ^(*t*+1)^*F*^⋆^, we calulate ^(*t*+1)^*V* ^*n*^ = ^(*t*+1)^*F*^*\**(*t*)^*S*^*n*^ and then update ^(*t*+1)^*G*^*n*^ for each modality.

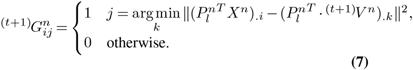

We then update ^(*t*+1)^*S*^*n*^ for each modality

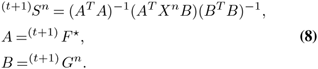

This iterative process allows us to refine the matrices and capture the evolving relationships between features and cells across multiple modalities.

#### A.4. Model selection strategy

After a certain number of iteration described in the previous section (also see Algorithm 1), denoted as *t*^⋆^, we have obtained an initial weighted shared matrix 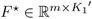, which provides preliminary insights into the cell clusters. Each row of *F*^⋆^ contains at most *K*_1_′ non-zero elements, and we can obtain cell cluster assignments for each cell by binarising *F*^⋆^:

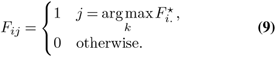

The cell-specific weights *w*^*n*^ for each modality indicate how each modality contributed to the reconstruction of the cell information. If a row of *F*^⋆^ has only one non-zero element, it signifies that each modality contributes shared information to the assignment of cell type for the corresponding cell. If a row of *F*^⋆^ has more than one non-zero elements, it suggests the presence of unique information from different modalities. In such cases, we select the cluster with the highest weight which has the highest relative importance.

##### Algorithm 1

TriTan Algorithm

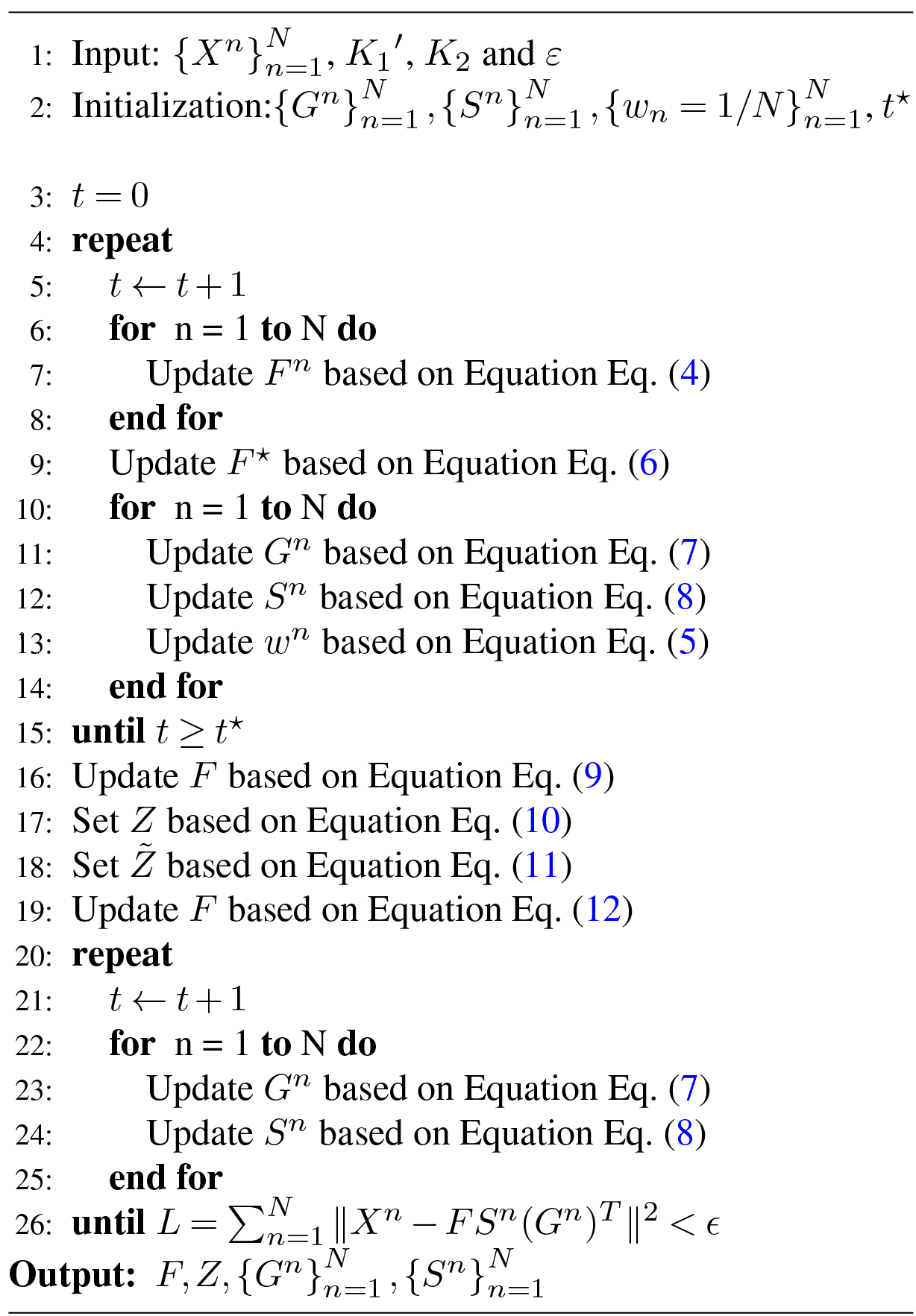

At the same time, we also build a shared cell embedding using our modality-specific weight, which effectively combines the contributions of each modality to capture the specific characteristics of individual cells:

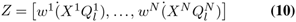

Then, we rearrange rows of *Z* into a row block matrix, denoted as 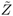, where the *i*-th block 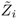 corresponds to cells being assigned to the *i*-th cluster, i.e.,

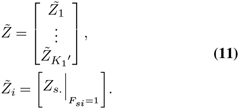

We next input 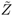 to HDBSCAN (15) for a local cell sub-type search. For each 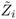 individually, HDBSCAN applies a density-based transformation to the data space, allowing it to identify stable dense points, which also leaves outliers that do not have any assigned cluster. We gather the centroids of each resulted clusters. Assuming in total *K*_1_ centres are obtained, we then concatenate them to form a new cluster centers matrix, denoted as *U*^*c*^. *U*^*c*^ is then used to update the shared cell cluster assignment matrix *F* as our final output:

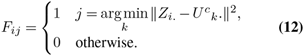

Clustering features can be generally more complex than clustering cells. So when it comes to determining the number of feature clusters, we adopt an exploratory approach. Here, we start with a large number of feature clusters and subsequently determine the appropriate number based on the condition of convergence as some of the clusters will be empty which we prune afterwards. This iterative process allows us to flexibly adapt to the complexity and structure of the data, ensuring accurate representation and meaningful interpretation of the feature clusters.

### B. Single-cell multi-modal datasets

We have used multiple publicly available single-cell multi-modal (two modalities - RNA and ATAC) datasets of varying sizes to comprehensively benchmark TriTan against other methods and leverage the output from TriTan for extensive downstream analyses. These include data generated using 10X Multiome from human peripheral blood mononuclear cells and bone marrow mononuclear cells (NeurIPS 2021, GSE194122) as well as mouse skin cells (GSM4156597) using SHARE-seq protocol (16). These data are summarised in Table 1.

**Table 1.** Multiomics datasets.

| Dataset | Protocol | Species | No. of cells | No. of cell types |
| --- | --- | --- | --- | --- |
| PBMC-10K | 10X Multiome | Human | 11787 | 13 |
| Skin-SHARE | SHARE-seq | Mouse | 34774 | 23 |
| NeurIPS 2021 | 10X Multiome | Human | 69249 | 22 |

The NeurIPS 2021 dataset (17) which is currently the largest dataset we used, was created as part of the NeurIPS competition and is utilised as a benchmark for accuracy and computational complexity of integration methods.

#### Data pre-processing

We performed uniform pre-processing for all the datasets using Scanpy pipeline for all the methods. For scRNA-seq matrices, we used `pp.highly_variable_genes()` to select the highly variable genes and then performed TF-IDF (Term Frequency-Inverse Document Frequency) transformation. For scATAC-seq modality, we first ran TF-IDF and then use `pp.highly_variable_genes()` to find the high variable peaks. We used different parameters of selecting highly variable features in regard to different datasets for scRNA-seq and scATAC-seq respectively. The relevant parameter values are shown in supplementary Table.S3.

### C. Implementation of competing methods

We selected three most popular and representative methods in single-cell multiomics data integration space. Below we describe these methods.

#### Seurat v4 (WNN)

The pipeline of Seurat v4 (2) uses Weighted Nearest Neighbor (WNN) approach for single-cell multiomics datasets. It consists three steps: firstly, for each modality, it pre-processes the dataset independently and reduces the data dimension. Then it learns cell-specific weights and constructs an integrated WNN graph. Finally, clustering and other downstream analysis and visualization are carried out based on the WNN graph. We followed the recommended tutorial from Seurat v4 website (https://satijalab.org/seurat/articles/multimodal_vignette.html) and executed Seurat v4 WNN in R for all datasets reported in this manuscript.

#### MOFA+

MOFA+ (4) is a popular method based on factor analysis that provides a general framework for the integration of multiomics datasets in an unsupervised manner. We followed the recommended tutorial from MOFA+ website (https://biofam.github.io/MOFA2/tutorials.html) and ran MOFA+ within Scanpy with default parameters for all datasets.

#### MOJITOO

MOJITOO (7) is a highly efficient method published recently which uses canonical correlation analysis to detect a shared representation for multiomics single-cell data. MOJITOO uses Seurat package to create a *SeuratObject* and reduces dimension for data in each modality. MOJITOO then implements CCA for an efficient and parameter-free identification of a shared representation of cells from multi-modal single-cell data. We ran MOJITOO for our applications using the tutorial (https://github.com/CostaLab/MOJITOO/) in R.

## Results

### D. Benchmarking of single-cell multi-modal integration methods

We evaluated TriTan and existing methods described above using three publicly available multi-modal datasets (Table 1) with two modalities.

Assessing the computational requirements of processing single-cell multiomics data with an ever-increasing number of cells is highly important metric in evaluating the effectiveness of methods. To do this, we randomly sampled cells from NeurIPS 2021 data which is our largest dataset and generated six datasets with an increasing number of cells, from 3000 to 60000. We evaluated TriTan and three methods described above using these six datasets and analysed the time and memory requirements in our benchmark by running them on a parallel 16-core job with a 512GB RAM/core node. For all methods, we used their default settings. Computational running time and memory comparisons are shown in Fig.2 where TriTan shows a very competitive performance overall. It is the fastest among all the methods, except in 60000 cells dataset with MOJITOO is slightly faster. However, WNN and MOJITOO, very similar to each other, still have the lowest computational demand for memory across all six datasets and TriTan ranks third.

**Fig. 2.**
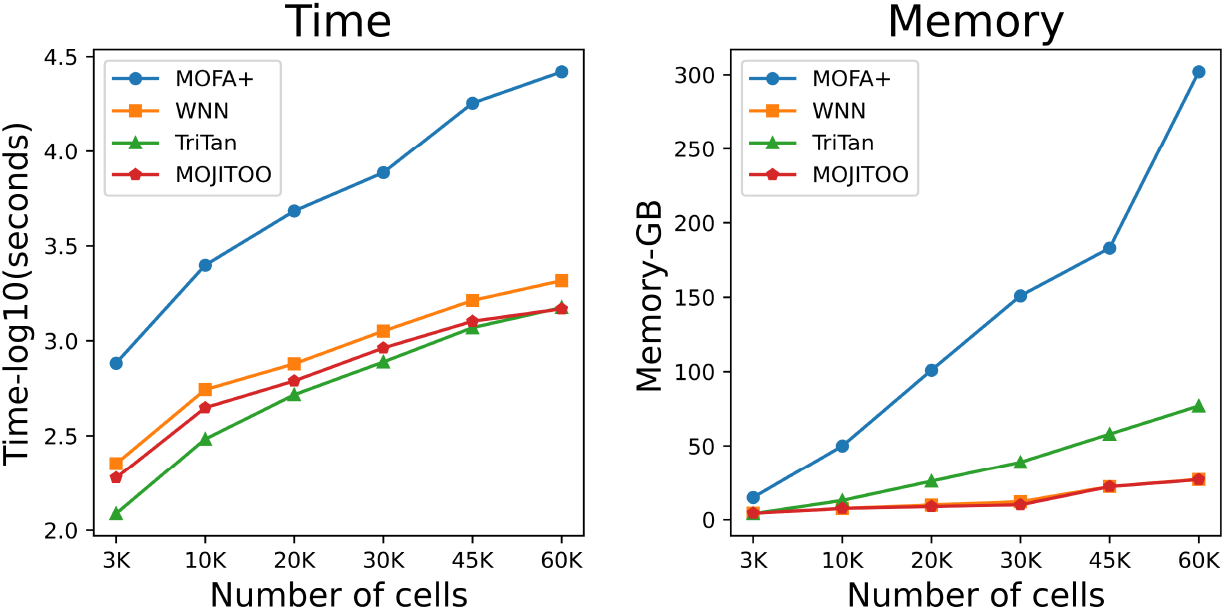
Time and memory usage for TriTan and three competing methods against different number of cells (x-axis) randomly sampled from he NeurIPS 2021 data, for each method, left subplot shows elapsed time (log10 of seconds) and right subplot shows peak memory (Gigabytes) required by each method.

Next, we assessed TriTan against other methods with regard to their ability to recover cell clusters using Adjusted Rand Index (ARI) (18) calculated by cell labels obtained from each method and the ground truth labels (Fig. 3). TriTan, WNN and MOFA+ can estimate the number of clusters and provide cell cluster labels, whereas MOJITOO only provides shared latent space. To obtain cell cluster labels for MOJITOO, we employed K-means clustering with the number of clusters based on ground truth which might give some advantage to MOJITOO in this comparison. Despite this, TriTan had the highest ARI scores in two out of three datasets, as shown in Fig. 3B.

**Fig. 3.**
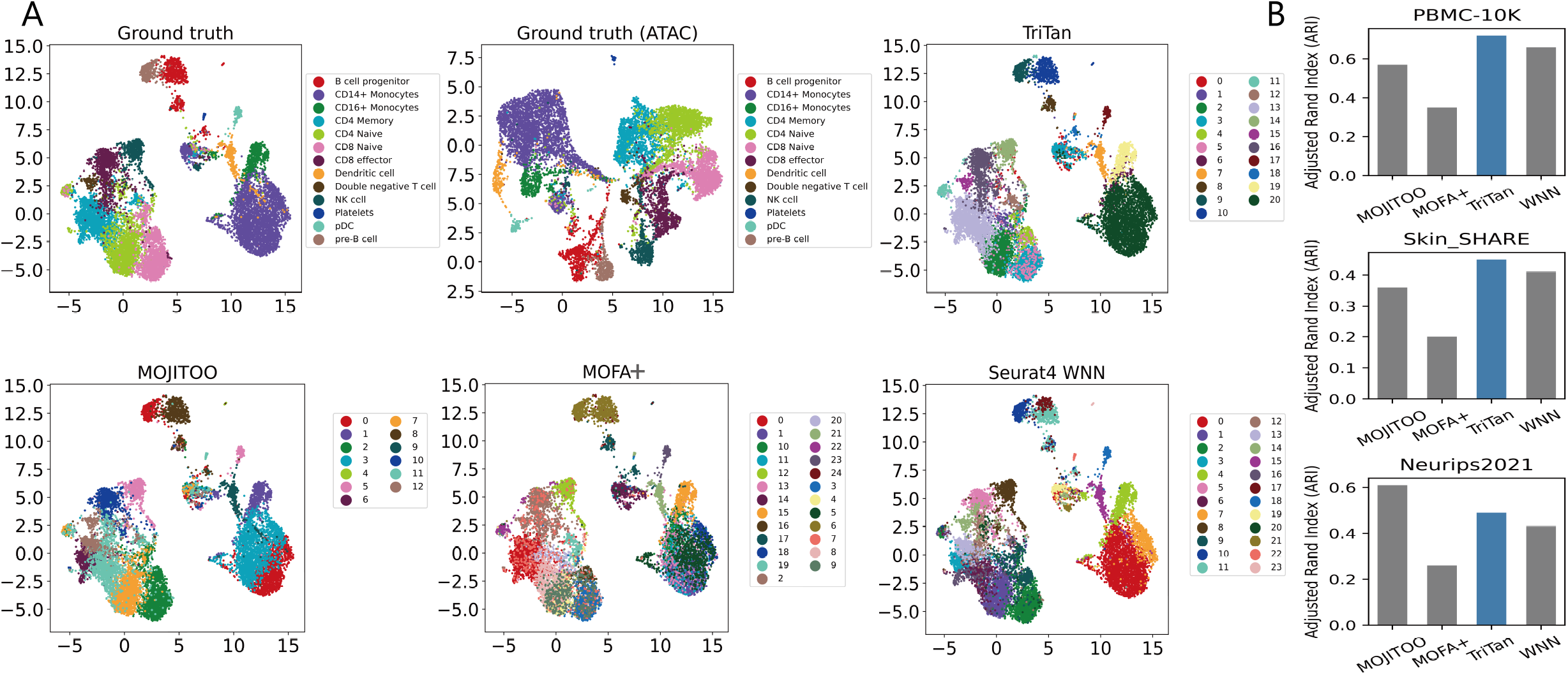
(**A**) UMAPs visualizations of cell clusters of 10X PBMC-10K data based on scRNA-seq data and scATAC-seq using ground truth labels as well as labels from MOFA+, WNN, MOJITOO, and TriTan. (**B**) Comparison of methods in terms of cell clustering accuracy (ARI) against ground truth labels for three different datasets.

Overall, this benchmarking analysis shows that TriTan is highly competitive against state-of-the-art methods in terms of accuracy in cell clustering as well as computational complexity. Furthermore, it provides additional advantages for downstream biological interpretation. As shown in next sections, it facilitates the association of cell clusters with feature clusters, as well as the prediction of relationships among features in different modalities.

### E. TriTan finds signature feature sets for each cell topic using association matrices

In this section, we examined the utility of association matrices produced by TriTan for different modalities. The use of these matrices provides additional information that is lacking in more commonly used two-matrix NMF methods. They enable the association of multiple feature clusters (gene sets and peak sets) with cell topics, as illustrated in Fig. 4A. The two heatmaps on the top display normalised association matrices derived from scATAC-seq and scRNA-seq data, while the bottom two heatmaps present binarised association matrices using a fixed threshold (we have chosen a strict threshold of 0.9 in this instance). These binarised matrices allow the discovery of gene sets and peak sets that define and differentiate different cell types. In our analysis, we observed that geneset-38, geneset-0, and geneset-28, along with peakset-13, peakset-18, and peakset-24, demonstrated strong association to plasmacytoid dendritic cells (pDCs). Additionally, geneset-9 and peakset-6 show a distinct association with double negative T cells, while peakset-26 (second row) exclusively identifies NK cells. It is worth noting that scATAc-seq data appears to play a more important role in defining double negative T cells and NK cells. Conversely, for other cell type like CD4 naive and CD8 naive cells, both signature gene sets and peak sets effectively distinguish between them.

**Fig. 4.**
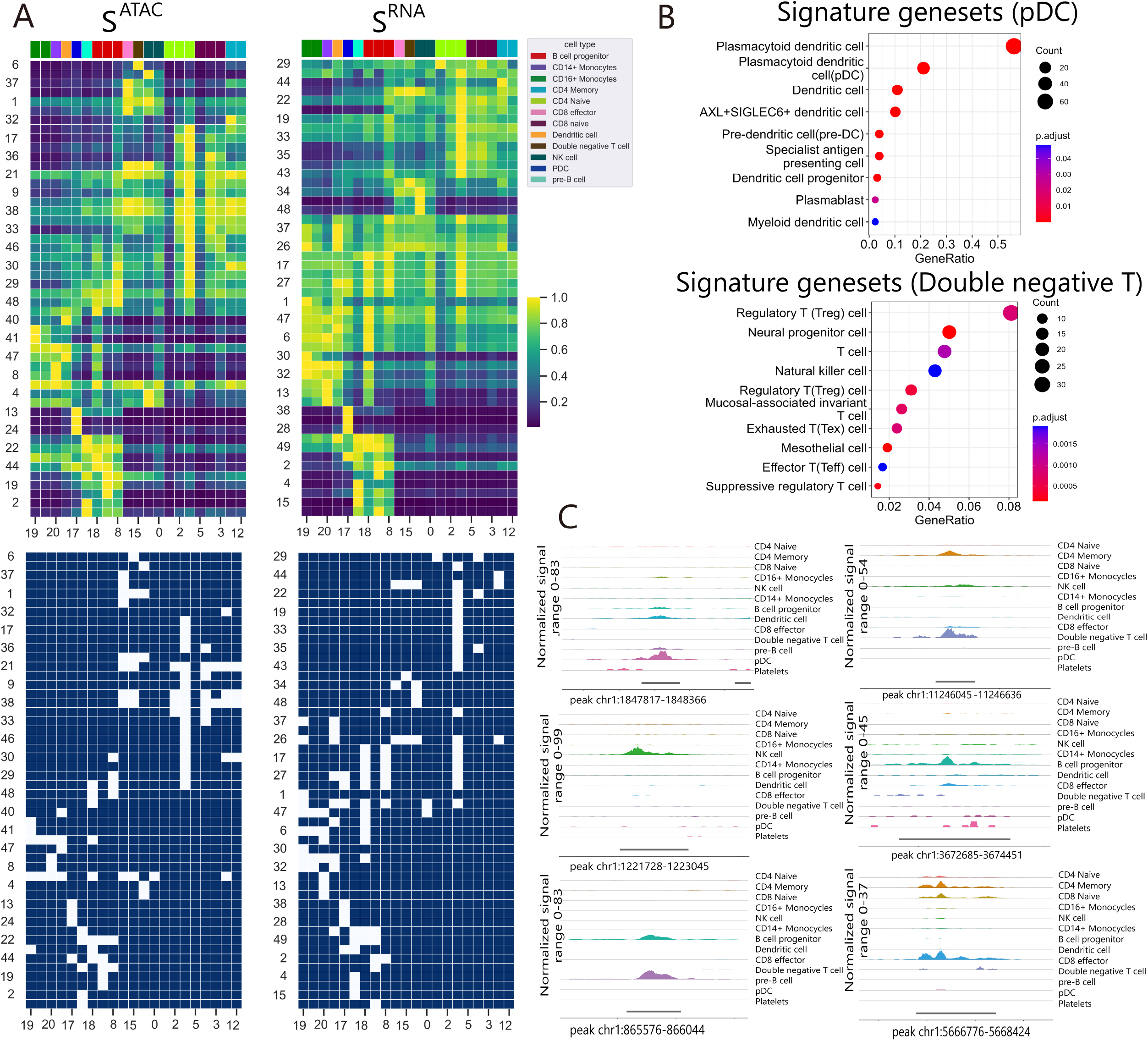
(**A**) Clustered heatmaps (top row) visualise normalised association matrices from each modality. Normalisation was done row-wise, dividing by the maximum value. For heatmaps, each row represents a feature cluster and each column represents a cell cluster. We also show two binarised association matrices with a fixed threshold (0.9) at the bottom so that we can define signature feature sets for each cell topic. Please note that, the heatmaps may not display the full labels on the X-axis and Y-axis. Complete heatmaps with full labels are shown in the supplementary Fig. S1. (**B**) Cell Marker over-representation analysis are performed with clusterProfiler 4.0 (19) using CellMarker 2.0 human dataset (20). Here we present the dotplots for signature gene sets of pDC cells and double negative T cells. More examples are shown in supplementary Fig. S4.**(C)** We selected one peak from the signature peak set for each cell topic and visualize DNA accessibility information using the `CoveragePlot()` function in Signac (21) for different cell types. We show six examples here for pDCs, Double negative T cells, NK cells, B cells progenitor, pre-B cells and CD8 effector cells. More examples are shown in supplementary Fig. S3.

In addition to identifying signature gene and peak sets with known cell types, TriTan can also identify potential subtypes within the ground truth, providing novel hypothesis for further investigation. For example, cell clusters 8, 10 and 18 detected by TriTan were all labelled as pro-B cell according to the ground truth; however, their gene expression levels and peak counts varied, suggesting potential differentiation or distinct stages within the pro-B cell population. Furthermore, our analysis has revealed that cell cluster 11 was predominantly characterised by geneset-29 (see supplementary Fig. S1 for full labels). It is noteworthy that this cluster exhibit a distinctive pattern that was observed exclusively in scRNA-seq data. Upon comparison with the ground truth labels, it appears that cluster 11 consisted of a combination of CD4 naive and CD8 naive cells.

To validate these findings, we used clusterProfiler 4.0 (19) to perform cell marker enrichment analysis on the pDC-specific gene sets and double negative T cell-specific gene sets. Cell-Marker 2.0 (20) human database provides a manually curated collection of experimentally supported markers of various cell types in different human tissues. These are presented in Fig.4B showing highly relevant terms enriched. Additionally, we showed coverage plots using selected peaks for each cell type as shown in Fig.4C. This approach allows us to visualise the enrichment level of individual peaks across different cell types and assess its specificity.

These results demonstrate TriTan’s ability to capture both modality-unique and modality-shared information and show that TriTan’s outputs, especially the middle association matrices, provide a unique advantage for characterising cell topics based on distinct combinations of gene sets and peak sets, enabling a comprehensive understanding of cellular processes and regulatory mechanisms.

### F. TriTan identifies Domains Of Regulatory Chromatin (DORCs)

For each cell topic, TriTan found signature gene sets and peak sets. Based on this, we aim to further explore gene-peak associations by utilising association matrices from RNA and ATAC modalities. For each gene set, Pearson correlations were computed against peak sets using their respective weights in association matrices across all cell topics. We visualised the correlation matrix using a clustered heatmap in Fig. 5A.

**Fig. 5.**
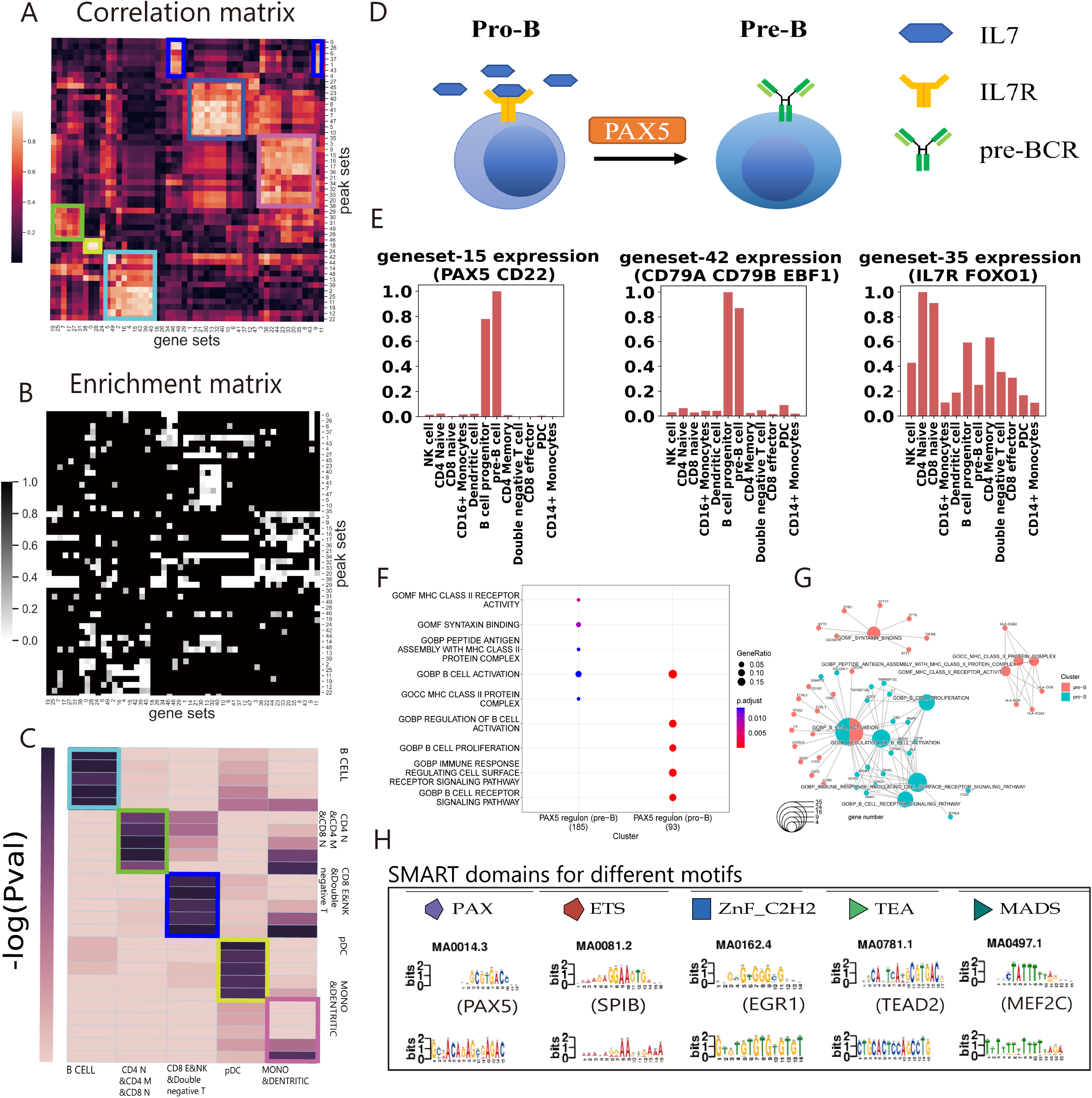
Downstream analysis of TriTan output. (**A**) The clustered heatmap of the correlation matrix, obtained from RNA and ATAC association matrices, shows clusters of gene sets and their potential regulating peak sets. This can be traced back to the cell type groups. (**B**) Clustered heatmap of the enrichment matrix from PEGS analysis, x-axis are the gene sets, and y-axis shows peak-sets (expanded to +/-2kb) with the same order of correlation matrix. (**C**) PEGS analysis for enrichment of signature gene sets (x-axis) and peak sets (y-axis, expanded to different genomic distances (2000,5000,10000,50000,150000)) derived from rectangles in (A). Axis labels show their corresponding cell type groups. The cell color represents -log10(p-value) of enrichment, and annotation of cells are the number of common genes for that particular combination of gene cluster (x-axis) and gene-set derived from the expanded peak-set (y-axis). (**D**) Cartoon diagram illustrating the transition of B cells from the pro-B stage to the pre-B stage which is regulated by PAX5 and the interaction between IL7 and IL7R. (**E**) The barplots showing some important gene sets’ expression reflected by their weights in association matrix across each cell type. (**F**) Comparative enrichment analysis was performed on PAX5 regulon (pre-B) and PAX5 regulon (pro-B). The regulons of PAX5 in pre-B and pro-B were analysed by GO enrichment, and the GO database used was from Molecular Signature Database (MSigDB) C5 (23, 24). The enrichment results were compared by the `compareCluster()` function of the clusterProfiler package, and visualised by `dotplot()` function. The size of the circle represents the gene ratio which is obtained by the number of genes belonging to the corresponding GO term compared to the total number of genes. The color represents the p-value. (**G**) Gene-Concept Network Plot of PAX5 regulon (pre-B) and PAX5 regulon (pro-B). The regulons of PAX5 in pre-B and pro-B were analysed by GO enrichment, and the GO database used was from Molecular Signature Database (MSigDB) C5 (23, 24). The enrichment results were compared by the `compareCluster()` function of the clusterProfiler package, and visualised by `centplot()` function. Centplot shows the relationship between genes and GO terms. Red represents genes from pre-B and corresponding enriched GO terms, and blue represents genes from pro-B and corresponding enriched GO terms. The size of the GO circle represents the number of enriched genes. (**H**) Five motifs correspond to five functional domains respectively. Enrichment analysis was performed on the 2kb region upstream and downstream of the TSS site of the gene from B cells identified by TriTan, in which the top 5 motifs were extracted and compared with the motifs in the Tomtom database, and only the interactions between motifs and transcription factors verified by ChIP were kept. The obtained transcription factors were put into the SMART (25) database for domain identification.

In order to test the mutual enrichment of identified gene and peak sets, we firstly used PEGS (Peak-set Enrichment of Gene-Sets)tool (22) and contrast the output against the correlation matrix obtained from scRNA and scATAC association matrices. Here, using PEGS, input peaks are extended to +/-2kb (promoter-proximal peaks) and the enrichment of the input gene set is calculated among the genes whose TSSs overlap with the extended peaks. We constructed the enrichment matrix that contains all the p-values for each gene set’s enrichment for each peak set as shown in Fig. 5B. The color of each cell in the heatmap represents the p-value of enrichment, and the x-axis and y-axis correspond to the gene sets and peak sets, respectively, in the same order as in the correlation matrix in Fig. 5A. Remarkably, the correlation matrix and the enrichment matrix exhibit a high degree of similarity, as shown in Fig. 5A&B, showing gene and peak sets with high degree of correlation and enrichment.

In the clustered heatmap of correlation matrix, the rectangles drawn using different colors represent gene sets and peak sets with high correlations and significant enrichment. Given our modality-specific association matrices, we observed that this structure corresponds to cell type groups, such as pro-B and pre-B cells group, CD14+ mono and CD16+ mono group, among others. Thus, we selected the signature gene sets and peak sets for these cell type groups and examined their enrichment using PEGS with varying peak extending distances (up to +/-150kb). As shown in Fig.5C, the combined signature gene sets of these clusters exhibited strong mutual enrichment to the combined signature peak sets. Based on this, we will further predict cell type specific regulons (TF and its target genes), as described in next section.

### G. TriTan predicts cell type-specific gene regulatory network and supports understanding of B cell trajectories at single-cell resolution

Here we used TriTan’s output to carry out downstream analysis in order to show its utility for finding cell type-specific regulatory networks. To demonstrate that the signature gene sets predicted by our method can improve the understanding of gene regulation, we performed motif analysis for signature peak sets of B cell populations from the PBMC-10K dataset. B lymphopoiesis, such as differentiation from pro-B cell (B-lymphoid progenitors) to pre-B cell (B cell precursors), is a genetically and epigenetically highly regulated process (26). Therefore, it is suitable for study on the transcription factor regulatory networks. The top five enriched motifs were identified. Using the Tomtom tool within MEME suite (27), we compared the identified motifs with those in the database and retained the ChIP-verified human transcription factors. The five motifs correspond to five functional domains respectively (see examples in Fig.5H).

- motif 1: mainly enriched to proteins with EST (E twenty-six) domains, such as SPIB, ELF1, and ETV1.
- motif 2: PAX (paired box) domain, such as PAX5.
- motif 3: TEA domain, such as TEAD1, TEAD2, and TEAD4.
- motif 4: MADS domain, such as MEF2A and MEF2C.
- motif 5: ZnF_C2H2 (Zinc finger C2H2-type) domain, such as EGR1 and KLF9.

These motif and domain pairs suggest various interactions between promoters and transcription factors during B cell development. SPIB, corresponding to the ETS domain and PU-box motif pair, inhibits human plasma cell differentiation by repressing BLIMP1 and XBP-1 expression (28), while the double mutation of its orthologs in mice develops pre-B cell acute lymphoblastic leukemia (29). MEF2C acts as a transcriptional activator of DNA repair to regulate the genome integrity and cell survival of pro-B, and its deficiency leads to a reduced chromatin accessible state in developmentally target regions in pre-B (30). TEAD1/2, a member of the Hippo signaling pathway, regulated the growth and self-renewal of non-lymphoid cells, was upregulated in leukemia, and was inhibited by IKAROS which regulates pre-B development (31). EGR1 has also been reported to be involved in the early differentiation process of B cells (32). PAX5 is a master regulator in the development of B cells (33). After losing the environment of IL7, pro-B cells lacking Pax5 can differentiate into other hematopoietic cell types, instead of entering B-cell lineage (34–36). As the Pax5 gene is activated, it can enter the pre-B stage, while differentiation from pre-B into plasma cells reduces the expression of Pax5 (33, 37). Furthermore, conditional deletion of Pax5 in pre-B resulted in the retrodifferentiation of cells into pro-B (33, 38).

Besides, differentially expressed features can also be found at the gene expression level. Unlike the traditional method of using an individual gene, TriTan can extract co-expressed gene sets through the similarity matrix. These gene sets not only enrich the genes that shared a similar function-related expression pattern but can more effectively and robustly identify differential expression events during cell development by calculating the relative expression level of the overall gene set. For example, in other types of non-B lymphocytes, the relative expression level of geneset-15 containing PAX5 is very low, but it is highly expressed in pro-B and pre-B, and the expression in pre-B is higher than that in pro-B (Fig.5E), which is consistent with the above. EBF1 is also a key regulatory factor for the early development of B cells, which specifically binds to the promoter of Cd79a (39, 40). Geneset-42 (Fig.5E) contains EBF1, CD79a, and CD79b, and its relative expression levels in pre-B and pro-B are much higher than those in other cell types, which well reflects the relationship and characteristics of the three genes in the early development of B cells. IL7R, the pro-B cell surface receptor, exists in geneset-35 (Fig.5E), and its relative expression level is significantly higher in pro-B cells than in pre-B cells. It is consistent with its function. Under the IL7 environment, IL7R activates the downstream JAK1/3 and STAT5a/b pathways, promotes the proliferation of pro-B, and prevents apoptosis and cell movement (41, 42). The pre-BCR signaling reduces the adhesion of pre-B by increasing CXCR4 and reducing FAK, which indirectly leads to reduced exposure of IL7 in pre-B cells and attenuates IL7R signaling (41, 42). In addition, the relatively high expression level of IL7R in T cells is also consistent with its previously reported role in T cell development and differentiation (41, 43). In this gene set, Bcl-2, which can be activated by IL7, and Foxo1, which acts as an enhancer to regulate the expression of IL7R, were also found (41, 44, 45). This indicates that our method enables the enrichment of gene sets with biological function correlation.

Differences in gene expression are regulated by transcription factors, so we next detected the difference in the PAX5 regulons between pro-B and pre-B (seen in supplementary Table.S1S2). To construct the PAX5 regulons specific to pro-B and pre-B cells, we used the signature gene sets identified by association matrices outputted by TriTan for each. We first used the Ensembl tool (46) to locate the transcription start site (TSS) of each gene and then selected those genes with PAX5 motifs enriched within 2kb upstream and down-stream of their TSS as the target genes for the PAX5 regulons. We performed comparative enrichment analysis (Seen in Fig.5F&G). The main enriched GO terms in pro-B are related to B cell proliferation, cell receptor signal transduction, and B cell activation, which may be related to IL7-mediated signal transduction (41, 43). The GO terms of B cell proliferation and cell receptor signal transduction are not enriched in pre-B, but there are specific terms such as MHCII and SYNTAXIN, which may be related to the production process of pre-BCR receptor in pre-B and the corresponding membrane fusion (47, 48).

Through this detailed analysis, we showed that the gene sets enriched by TriTan reflect the changes in gene expression during B cell development, and the regulon identified based on TriTan can also be functionally related, suggesting its potential in the study of regulatory mechanisms of cell types.

## Conclusion

Here, we have presented TriTan, a triple non-negative matrix factorisation algorithm for integrative analysis of single cell multiomics data. TriTan is highly competitive against three state-of-the-art methods in terms of computational complexity and accuracy in terms of clustering within our benchmark study using three single-cell multiomics public datasets from two different experimental technologies. It is important to note that clustering evaluation metric (ARI scores) requires ground truth labels, which may be biased towards the methods used to derive those labels, such as WNN. Additionally, in our evaluation of MOJITOO, given that it does not provide any model selection strategy, we used K-means to cluster cells into the same number of clusters as the ground truth, which may also introduce bias in the favour of MOJITOO. Importantly, we want to highlight the biological interpretation TriTan can provide to support downstream analysis. TriTan provides a unique capability for the identification of cell type specific signature feature sets through our middle association matrices. These matrices reflect the weights of each feature set across different cell types, enabling users to explore how feature sets from different modalities identify or co-identify a cell type. By computing the correlation between feature sets from different modalities, gene-peak linkages and domains of regulatory chromatin (DORCs) can be identified. Finally, by scanning the motifs over the specific DORCs for each cell type, we enable the prediction of cell type specific regulons and gene regulation networks. This powerful feature of TriTan will enable researchers to gain deeper insights into the complex interactions between different molecular features. TriTan is available as a Python package, accompanied with easy-to-use Jupyter notebook and example data.

## Data availability

The data underlying this article are available at https://doi.org/10.48420/23283998.v1 and https://doi.org/10.48420/23289797.v1.

## References

1. Mirjana Efremova and Sarah A Teichmann. Computational methods for single-cell omics across modalities. Nature methods, 17(1):14–17, 2020.

2. Yuhan Hao, Stephanie Hao, Erica Andersen-Nissen, William M Mauck III, Shiwei Zheng, Andrew Butler, Maddie J Lee, Aaron J Wilk, Charlotte Darby, Michael Zager, et al. Integrated analysis of multimodal single-cell data. Cell, 184(13):3573–3587, 2021.

3. Rohit Singh, Brian L Hie, Ashwin Narayan, and Bonnie Berger. Schema: metric learning enables interpretable synthesis of heterogeneous single-cell modalities. Genome biology, 22(1):1–24, 2021.

4. Ricard Argelaguet, Damien Arnol, Danila Bredikhin, Yonatan Deloro, Britta Velten, John C Marioni, and Oliver Stegle. Mofa+: a statistical framework for comprehensive integration of multi-modal single-cell data. Genome biology, 21(1):1–17, 2020.

5. Suoqin Jin, Lihua Zhang, and Qing Nie. scai: an unsupervised approach for the integrative analysis of parallel single-cell transcriptomic and epigenomic profiles. Genome biology, 21 (1):1–19, 2020.

6. April R Kriebel and Joshua D Welch. Uinmf performs mosaic integration of single-cell multiomic datasets using nonnegative matrix factorization. Nature communications, 13(1):1–17, 2022.

7. Mingbo Cheng, Zhijian Li, and Ivan Gesteira Costa Filho. Mojitoo: a fast and universal method for integration of multimodal single cell data. bioRxiv, 2022.

8. Adam Gayoso, Zoë Steier, Romain Lopez, Jeffrey Regier, Kristopher L Nazor, Aaron Streets, and Nir Yosef. Joint probabilistic modeling of single-cell multi-omic data with totalvi. Nature methods, 18(3):272–282, 2021.

9. Xinming Tu, Zhi-Jie Cao, Chen-Rui Xia, Sara Mostafavi, and Ge Gao. Cross-linked unified embedding for cross-modality representation learning. In Alice H. Oh, Alekh Agarwal, Danielle Belgrave, and Kyunghyun Cho, editors, Advances in Neural Information Processing Systems, 2022.

10. Danila Bredikhin, Ilia Kats, and Oliver Stegle. Muon: multimodal omics analysis framework. Genome Biology, 23(1):42, 2022.

11. F Alexander Wolf, Philipp Angerer, and Fabian J Theis. Scanpy: large-scale single-cell gene expression data analysis. Genome biology, 19:1–5, 2018.

12. Chris Ding, Tao Li, Wei Peng, and Haesun Park. Orthogonal nonnegative matrix tfactorizations for clustering. In Proceedings of the 12th ACM SIGKDD international conference on Knowledge discovery and data mining, pages 126–135, 2006.

13. Daniel Lee and H Sebastian Seung. Algorithms for non-negative matrix factorization. Advances in neural information processing systems, 13, 2000.

14. Antonella Falini. A review on the selection criteria for the truncated svd in data science applications. Journal of Computational Mathematics and Data Science, 5:100064, 2022. ISSN 2772-4158. doi: https://doi.org/10.1016/j.jcmds.2022.100064.

15. Leland McInnes, John Healy, and Steve Astels. hdbscan: Hierarchical density based clustering. The Journal of Open Source Software, 2(11), mar 2017. doi: 10.21105/joss.00205.

16. Sai Ma, Bing Zhang, Lindsay M. LaFave, Andrew S. Earl, Zachary Chiang, Yan Hu, Jiarui Ding, Alison Brack, Vinay K. Kartha, Tristan Tay, Travis Law, Caleb Lareau, Ya-Chieh Hsu, Aviv Regev, and Jason D. Buenrostro. Chromatin potential identified by shared single-cell profiling of rna and chromatin. Cell, 183(4):1103–1116.e20, 2020. ISSN 0092-8674. doi: https://doi.org/10.1016/j.cell.2020.09.056.

17. Malte Luecken, Daniel Burkhardt, Robrecht Cannoodt, Christopher Lance, Aditi Agrawal, Hananeh Aliee, Ann Chen, Louise Deconinck, Angela Detweiler, Alejandro Granados, Shelly Huynh, Laura Isacco, Yang Kim, Dominik Klein, BONY DE KUMAR, Sunil Kuppasani, Heiko Lickert, Aaron McGeever, Joaquin Melgarejo, Honey Mekonen, Maurizio Morri, Michaela Müller, Norma Neff, Sheryl Paul, Bastian Rieck, Kaylie Schneider, Scott Steelman, Michael Sterr, Daniel Treacy, Alexander Tong, Alexandra-Chloe Villani, Guilin Wang, Jia Yan, Ce Zhang, Angela Pisco, Smita Krishnaswamy, Fabian Theis, and Jonathan M Bloom. A sandbox for prediction and integration of dna, rna, and proteins in single cells. In J. Vanschoren and S. Yeung, editors, Proceedings of the Neural Information Processing Systems Track on Datasets and Benchmarks, volume 1. Curran, 2021.

18. Lawrence Hubert and Phipps Arabie. Comparing partitions. Journal of classification, 2: 193–218, 1985.

19. Tianzhi Wu, Erqiang Hu, Shuangbin Xu, Meijun Chen, Pingfan Guo, Zehan Dai, Tingze Feng, Lang Zhou, Wenli Tang, Li Zhan, et al. clusterprofiler 4.0: A universal enrichment tool for interpreting omics data. The Innovation, 2(3):100141, 2021.

20. Congxue Hu, Tengyue Li, Yingqi Xu, Xinxin Zhang, Feng Li, Jing Bai, Jing Chen, Wenqi Jiang, Kaiyue Yang, Qi Ou, Xia Li, Peng Wang, and Yunpeng Zhang. CellMarker 2.0: an updated database of manually curated cell markers in human/mouse and web tools based on scRNA-seq data. Nucleic Acids Research, 51(D1):D870–D876,10 2022. ISSN 0305-1048. doi: 10.1093/nar/gkac947.

21. Tim Stuart, Avi Srivastava, Shaista Madad, Caleb Lareau, and Rahul Satija. Single-cell chromatin state analysis with signac. Nature Methods, 2021. doi: 10.1038/s41592-021-01282-5.

22. Yang Sh et al. Briggs P, Hunter AL. Pegs: An efficient tool for gene set enrichment within defined sets of genomic intervals. F1000Research 2021, 2(3):100141, 2021.

23. Arthur Liberzon, Aravind Subramanian, Reid Pinchback, Helga Thorvaldsdóttir, Pablo Tamayo, and Jill P. Mesirov. Molecular signatures database (MSigDB) 3.0. 27(12):1739–1740, 2011. ISSN 1367-4803. doi: 10.1093/bioinformatics/btr260.

24. Aravind Subramanian, Pablo Tamayo, Vamsi K. Mootha, Sayan Mukherjee, Benjamin L. Ebert, Michael A. Gillette, Amanda Paulovich, Scott L. Pomeroy, Todd R. Golub, Eric S. Lander, and Jill P. Mesirov. Gene set enrichment analysis: A knowledge-based approach for interpreting genome-wide expression profiles. 102(43):15545–15550, 2005. doi: 10.1073/pnas.0506580102. Publisher: Proceedings of the National Academy of Sciences.

25. Ivica Letunic, Supriya Khedkar, and Peer Bork. SMART: recent updates, new developments and status in 2020. Nucleic Acids Research, 49(D1):D458–D460, 10 2020. ISSN 0305-1048. doi: 10.1093/nar/gkaa937.

26. Sören Boller and Rudolf Grosschedl. The regulatory network of B-cell differentiation: A focused view of early B-cell factor 1 function. Immunological Reviews, 261(1):102–115, September 2014. ISSN 0105-2896. doi: 10.1111/imr.12206.

27. Timothy L. Bailey, Mikael Boden, Fabian A. Buske, Martin Frith, Charles E. Grant, Luca Clementi, Jingyuan Ren, Wilfred W. Li, and William S. Noble. MEME Suite: tools for motif discovery and searching. Nucleic Acids Research, 37(suppl_2):W202–W208, 05 2009. ISSN 0305-1048. doi: 10.1093/nar/gkp335.

28. Heike Schmidlin, Sean A. Diehl, Maho Nagasawa, Ferenc A. Scheeren, Remko Schotte, Christel H. Uittenbogaart, Hergen Spits, and Bianca Blom. Spi-B inhibits human plasma cell differentiation by repressing BLIMP1 and XBP-1 expression. Blood, 112(5):1804–1812, September 2008. ISSN 1528-0020. doi: 10.1182/blood-2008-01-136440.

29. Kristen M. Sokalski, Stephen K. H. Li, Ian Welch, Heather-Anne T. Cadieux-Pitre, Marek R. Gruca, and Rodney P. DeKoter. Deletion of genes encoding PU.1 and Spi-B in B cells impairs differentiation and induces pre-B cell acute lymphoblastic leukemia. Blood, 118 (10):2801–2808, September 2011. ISSN 1528-0020. doi: 10.1182/blood-2011-02-335539.

30. Wenyuan Wang, Tonis Org, Amélie Montel-Hagen, Peter D. Pioli, Dan Duan, Edo Israely, Daniel Malkin, Trent Su, Johanna Flach, Siavash K. Kurdistani, Robert H. Schiestl, and Hanna K. A. Mikkola. MEF2C protects bone marrow B-lymphoid progenitors during stress haematopoiesis. Nature Communications, 7:12376, August 2016. ISSN 2041-1723. doi: 10.1038/ncomms12376.

31. Yeguang Hu, Zhihong Zhang, Mariko Kashiwagi, Toshimi Yoshida, Ila Joshi, Nilamani Jena, Rajesh Somasundaram, Akinola Olumide Emmanuel, Mikael Sigvardsson, Julien Fitamant, Nabeel El-Bardeesy, Fotini Gounari, Richard A. Van Etten, and Katia Georgopoulos. Superenhancer reprogramming drives a B-cell–epithelial transition and high-risk leukemia. Genes & Development, 30(17):1971–1990, September 2016. ISSN 0890-9369. doi: 10.1101/gad.283762.116.

32. A. Dinkel, K. Warnatz, B. Ledermann, A. Rolink, P. F. Zipfel, K. Bürki, and H. Eibel. The transcription factor early growth response 1 (Egr-1) advances differentiation of pre-B and immature B cells. The Journal of Experimental Medicine, 188(12):2215–2224, December 1998. ISSN 0022-1007. doi: 10.1084/jem.188.12.2215.

33. César Cobaleda, Alexandra Schebesta, Alessio Delogu, and Meinrad Busslinger. Pax5: The guardian of B cell identity and function. Nature Immunology, 8(5):463–470, May 2007. ISSN 1529-2908. doi: 10.1038/ni1454.

34. S. L. Nutt, B. Heavey, A. G. Rolink, and M. Busslinger. Commitment to the B-lymphoid lineage depends on the transcription factor Pax5. Nature, 401(6753):556–562, October 1999. ISSN 0028-0836. doi: 10.1038/44076.

35. A. G. Rolink, S. L. Nutt, F. Melchers, and M. Busslinger. Long-term in vivo reconstitution of T-cell development by Pax5-deficient B-cell progenitors. Nature, 401(6753):603–606, October 1999. ISSN 0028-0836. doi: 10.1038/44164.

36. Christoph Schaniel, Ludovica Bruno, Fritz Melchers, and Antonius G. Rolink. Multiple hematopoietic cell lineages develop in vivo from transplanted Pax5-deficient pre-B I-cell clones. Blood, 99(2):472–478, January 2002. ISSN 0006-4971. doi: 10.1182/blood.v99.2.472.

37. Meinrad Busslinger. Transcriptional control of early B cell development. Annual Review of Immunology, 22:55–79, 2004. ISSN 0732-0582. doi: 10.1146/annurev.immunol.22.012703.104807.

38. Alessio Delogu, Alexandra Schebesta, Qiong Sun, Katharina Aschenbrenner, Thomas Perlot, and Meinrad Busslinger. Gene repression by Pax5 in B cells is essential for blood cell homeostasis and is reversed in plasma cells. Immunity, 24(3):269–281, March 2006. ISSN 1074-7613. doi: 10.1016/j.immuni.2006.01.012.

39. Sören Boller, Rui Li, and Rudolf Grosschedl. Defining B Cell Chromatin: Lessons from EBF1. Trends in Genetics, 34(4):257–269, April 2018. ISSN 0168-9525. doi: 10.1016/j.tig.2017.12.014.

40. Rui Li, Pierre Cauchy, Senthilkumar Ramamoorthy, Sören Boller, Lukas Chavez, and Rudolf Grosschedl. Dynamic EBF1 occupancy directs sequential epigenetic and transcriptional events in B-cell programming. Genes & Development, 32(2):96–111, January 2018. ISSN 1549-5477. doi: 10.1101/gad.309583.117.

41. João T. Barata, Scott K. Durum, and Benedict Seddon. Flip the coin: IL-7 and IL-7R in health and disease. Nature Immunology, 20(12):1584–1593, December 2019. ISSN 1529-2916. doi: 10.1038/s41590-019-0479-x.

42. Chris Fistonich, Sandra Zehentmeier, Jeffrey J. Bednarski, Runfeng Miao, Hilde Schjerven, Barry P. Sleckman, and João P. Pereira. Cell circuits between B cell progenitors and IL-7+ mesenchymal progenitor cells control B cell development. The Journal of Experimental Medicine, 215(10):2586–2599, October 2018. ISSN 1540-9538. doi: 10.1084/jem.20180778.

43. Eva Buentke, Anne Mathiot, Mauro Tolaini, James Di Santo, Rose Zamoyska, and Benedict Seddon. Do CD8 effector cells need IL-7R expression to become resting memory cells? Blood, 108(6):1949–1956, September 2006. ISSN 0006-4971. doi: 10.1182/blood-2006-04-016857.

44. Yann M. Kerdiles, Daniel R. Beisner, Roberto Tinoco, Anne S. Dejean, Diego H. Castrillon, Ronald A. DePinho, and Stephen M. Hedrick. Foxo1 links homing and survival of naive T cells by regulating L-selectin, CCR7 and interleukin 7 receptor. Nature Immunology, 10(2):176–184, February 2009. ISSN 1529-2916. doi: 10.1038/ni.1689.

45. Daniel Ribeiro, Alice Melão, Ruben van Boxtel, Cristina I. Santos, Ana Silva, Milene C. Silva, Bruno A. Cardoso, Paul J. Coffer, and João T. Barata. STAT5 is essential for IL-7-mediated viability, growth, and proliferation of T-cell acute lymphoblastic leukemia cells. Blood Advances, 2(17):2199–2213, September 2018. ISSN 2473-9537. doi: 10.1182/bloodadvances.2018021063.

46. Fiona Cunningham, James E Allen, Jamie Allen, Jorge Alvarez-Jarreta, M Ridwan Amode, Irina M Armean, Olanrewaju Austine-Orimoloye, Andrey G Azov, If Barnes, Ruth Bennett, Andrew Berry, Jyothish Bhai, Alexandra Bignell, Konstantinos Billis, Sanjay Boddu, Lucy Brooks, Mehrnaz Charkhchi, Carla Cummins, Luca Da Rin Fioretto, Claire Davidson, Kamalkumar Dodiya, Sarah Donaldson, Bilal El Houdaigui, Tamara El Naboulsi, Reham Fatima, Carlos Garcia Giron, Thiago Genez, Jose Gonzalez Martinez, Cristina Guijarro-Clarke, Arthur Gymer, Matthew Hardy, Zoe Hollis, Thibaut Hourlier, Toby Hunt, Thomas Juettemann, Vinay Kaikala, Mike Kay, Ilias Lavidas, Tuan Le, Diana Lemos, José Carlos Marugán, Shamika Mohanan, Aleena Mushtaq, Marc Naven, Denye N Ogeh, Anne Parker, Andrew Parton, Malcolm Perry, Ivana PiliŽota, Irina Prosovetskaia, Manoj Pandian Sakthivel, Ahamed Imran Abdul Salam, Bianca M Schmitt, Helen Schuilenburg, Dan Shep-pard, José G Pérez-Silva, William Stark, Emily Steed, Kyösti Sutinen, Ranjit Sukumaran, Dulika Sumathipala, Marie-Marthe Suner, Michal Szpak, Anja Thormann, Francesca Floriana Tricomi, David Urbina-Gómez, Andres Veidenberg, Thomas A Walsh, Brandon Walts, Natalie Willhoft, Andrea Winterbottom, Elizabeth Wass, Marc Chakiachvili, Bethany Flint, Adam Frankish, Stefano Giorgetti, Leanne Haggerty, Sarah E Hunt, Garth R IIsley, Jane E Loveland, Fergal J Martin, Benjamin Moore, Jonathan M Mudge, Matthieu Muffato, Emily Perry, Magali Ruffier, John Tate, David Thybert, Stephen J Trevanion, Sarah Dyer, Peter W Harrison, Kevin L Howe, Andrew D Yates, Daniel R Zerbino, and Paul Flicek. Ensembl 2022. Nucleic Acids Research, 50(D1):D988–D995, 11 2021. ISSN 0305-1048. doi: 10.1093/nar/gkab1049.

47. Mark K. Bennett, JoséE. Garcia-Arrarás, Lisa A. Elferink, Karen Peterson, Anne M. Fleming, Christopher D. Hazuka, and Richard H. Scheller. The syntaxin family of vesicular transport receptors. Cell, 74(5):863–873, September 1993. ISSN 0092-8674. doi: 10.1016/0092-8674(93)90466-4.

48. Julia Merkenschlager, Urszula Eksmond, Luca Danelli, Jan Attig, George R. Young, Carla Nowosad, Pavel Tolar, and George Kassiotis. MHC class II cell-autonomously regulates self-renewal and differentiation of normal and malignant B cells. Blood, 133(10):1108–1118, March 2019. ISSN 0006-4971. doi: 10.1182/blood-2018-11-885467.

