## supplementary for "TriTan: An efficient triple non-negative matrix factorisation method for integrative analysis of single-cell multiomics data"

**Fig. S1**: Clustered heatmaps visualizing normalized association matrices for each modality with full labels.

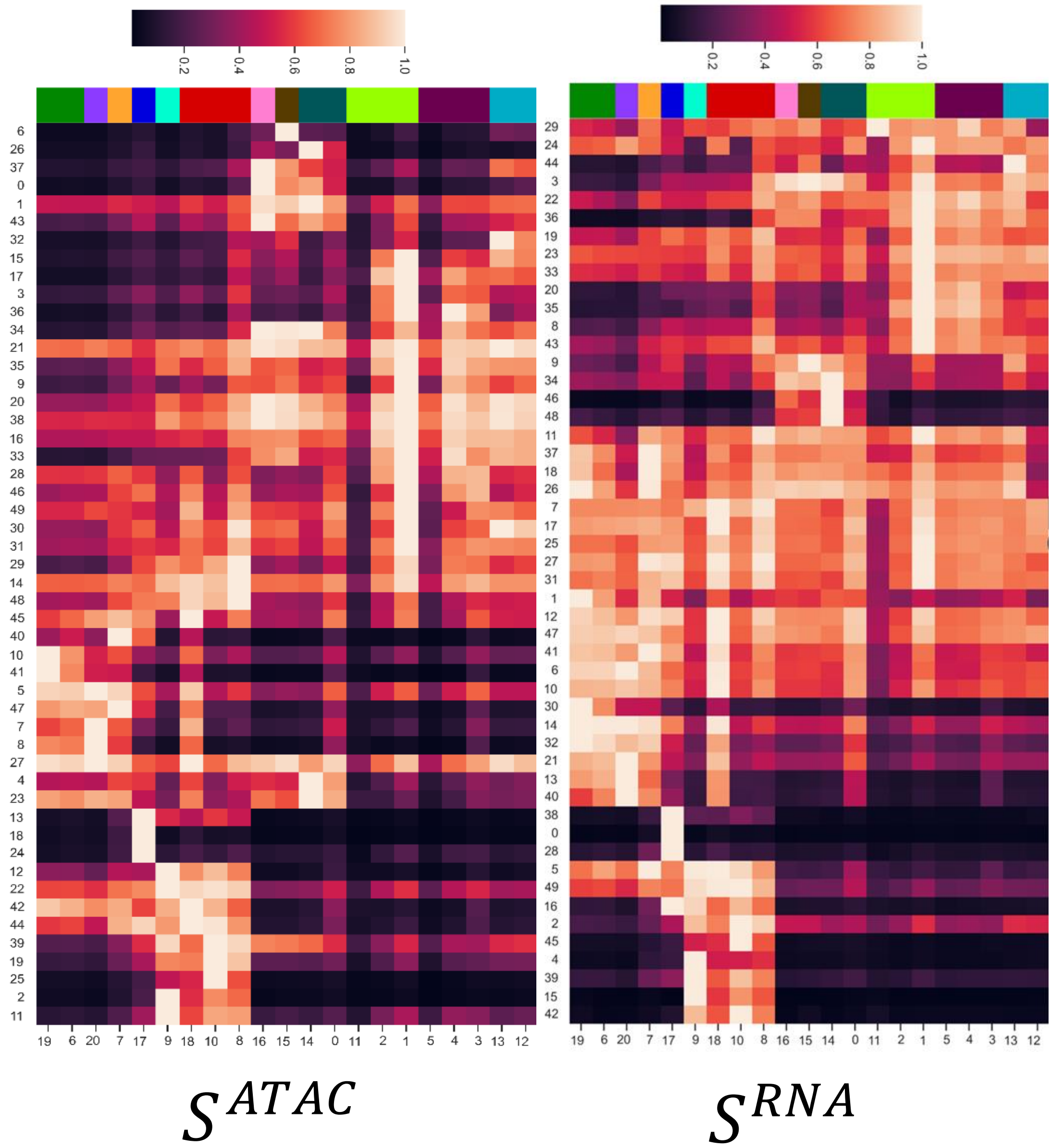

**Fig. S2:** UMAPs visualizations of cell clusters of 10X PBMC-10K data based on scRNA-seq data and scATAC-seq using ground truth labels as well as labels from MOFA+, WNN, MOJITOO, and TriTan.

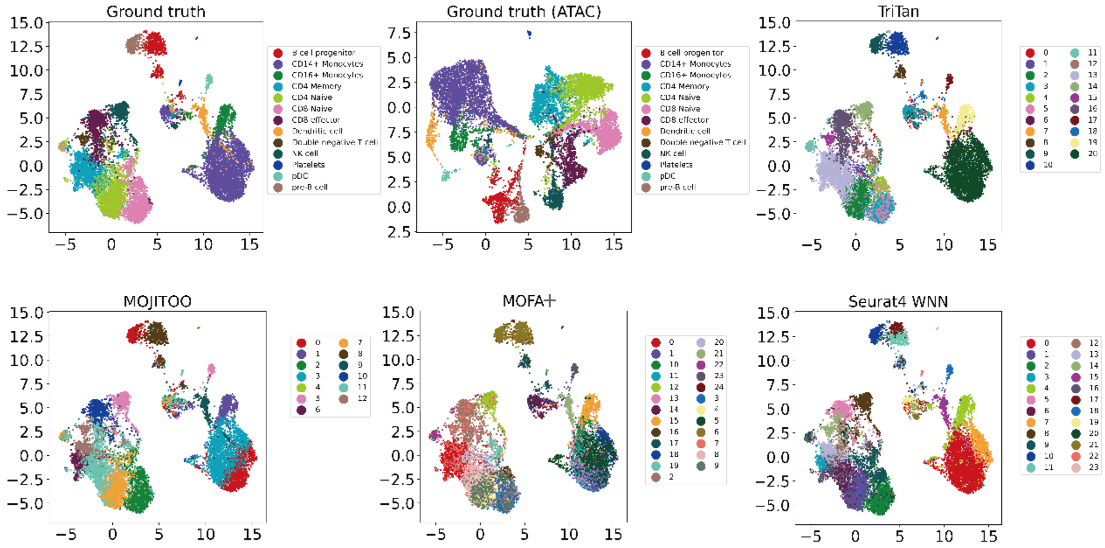

**Fig. S3:** Using a selected peak from the signature peak set for each cell topic to visualize DNA accessibility information using the CoveragePlot() function in Signac for CD14+ Mono, CD4 Naive, CD4 Memory, CD16+ Mono, and Dendritic cells.

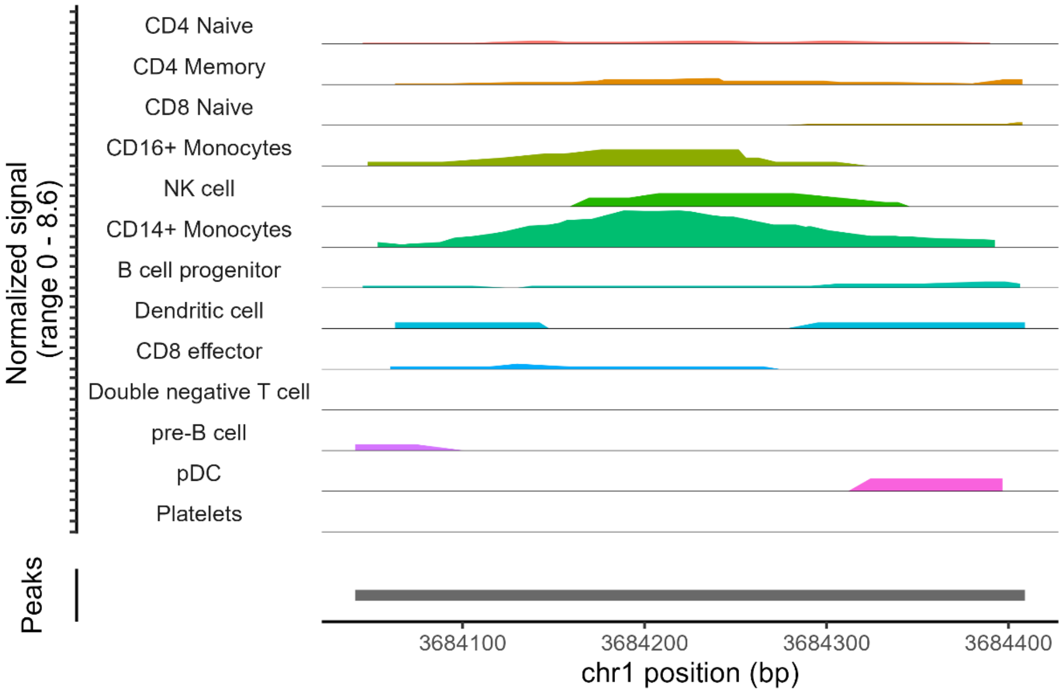


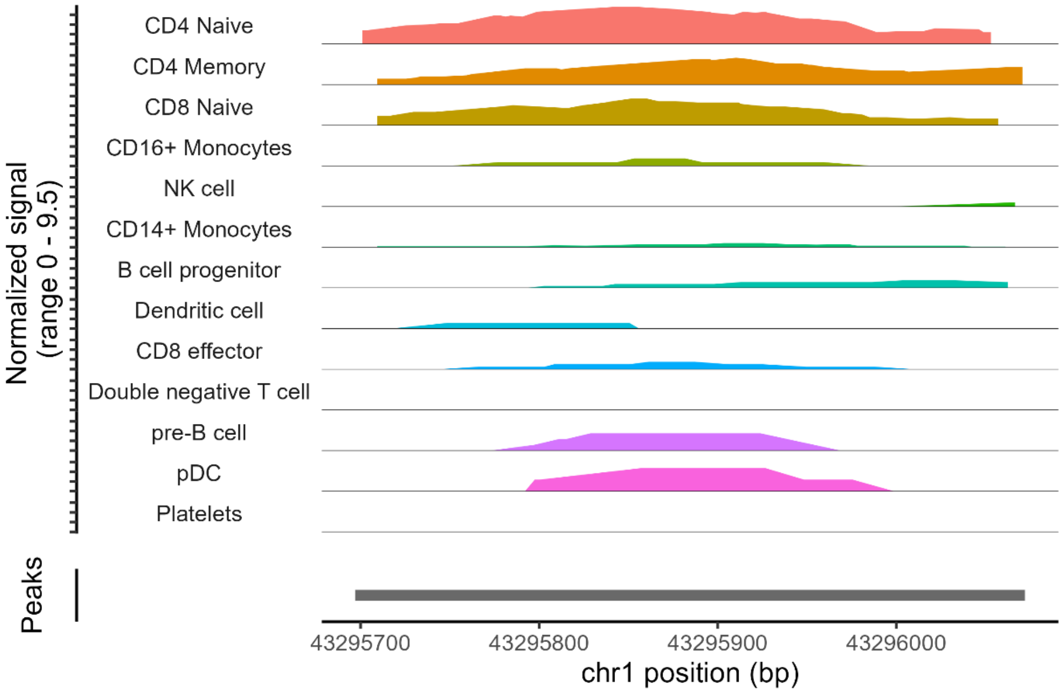

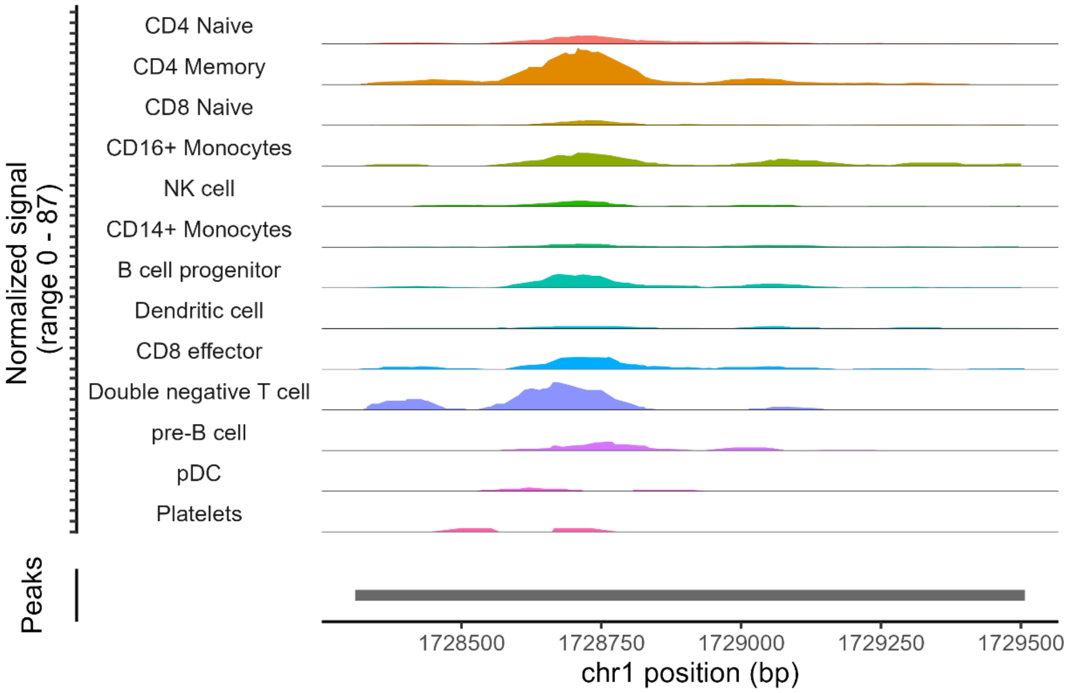
Normalized signal
(range 0 - 87)
CD4 Naive
CD4 Memory
CD8 Naive
CD16+ Monocytes
NK cell
CD14+ Monocytes
B cell progenitor
Dendritic cell
CD8 effector
Double negative T cell
pre-B cell
pDC
Platelets
Peaks
1728500
1728750
1729000
1729250
1729500
chr1 position (bp)

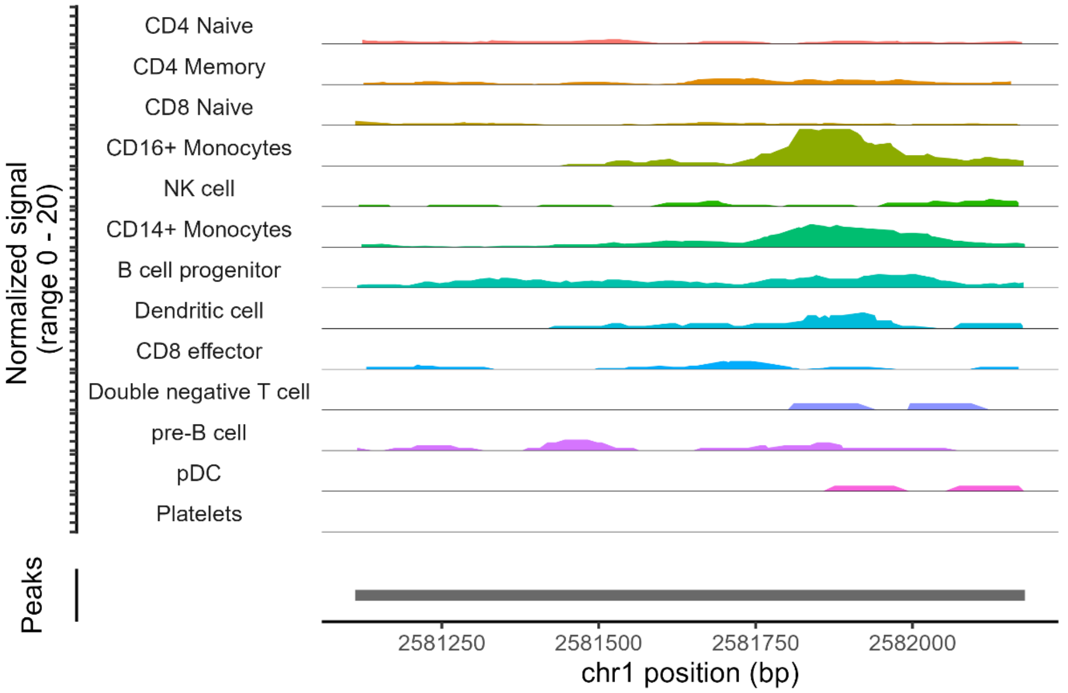
Normalized signal
(range 0 - 20)
CD4 Naive
CD4 Memory
CD8 Naive
CD16+ Monocytes
NK cell
CD14+ Monocytes
B cell progenitor
Dendritic cell
CD8 effector
Double negative T cell
pre-B cell
pDC
Platelets
Peaks
2581250
2581500
2581750
2582000
chr1 position (bp)

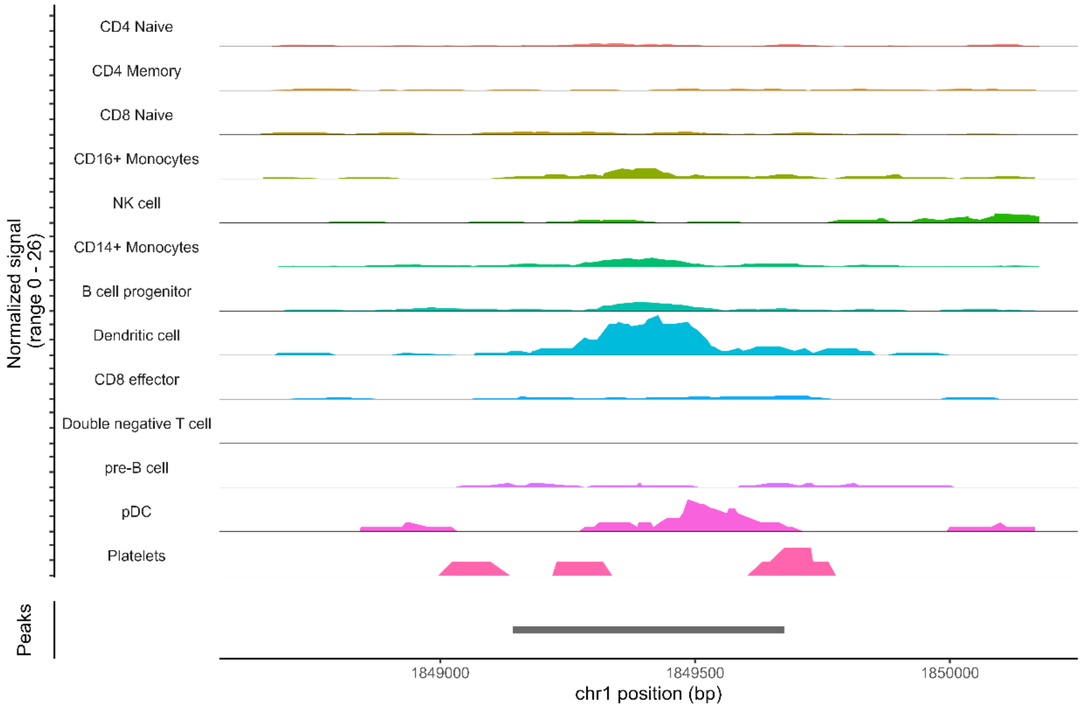
Normalized signal
(range 0 - 26)
CD4 Naive
CD4 Memory
CD8 Naive
CD16+ Monocytes
NK cell
CD14+ Monocytes
B cell progenitor
Dendritic cell
CD8 effector
Double negative T cell
pre-B cell
pDC
Platelets
Peaks
1849000
1849500
1850000
chr1 position (bp)

**Fig. S4:** Cell Marker over-representation analysis performed by clusterProfiler 4.0 using CellMarker 2.0 human dataset. Here the dotplots are drawn for specific gene sets of a) NK cells, b) CD16+ Mono, c) CD14+ Mono, and d) T cells.

a)

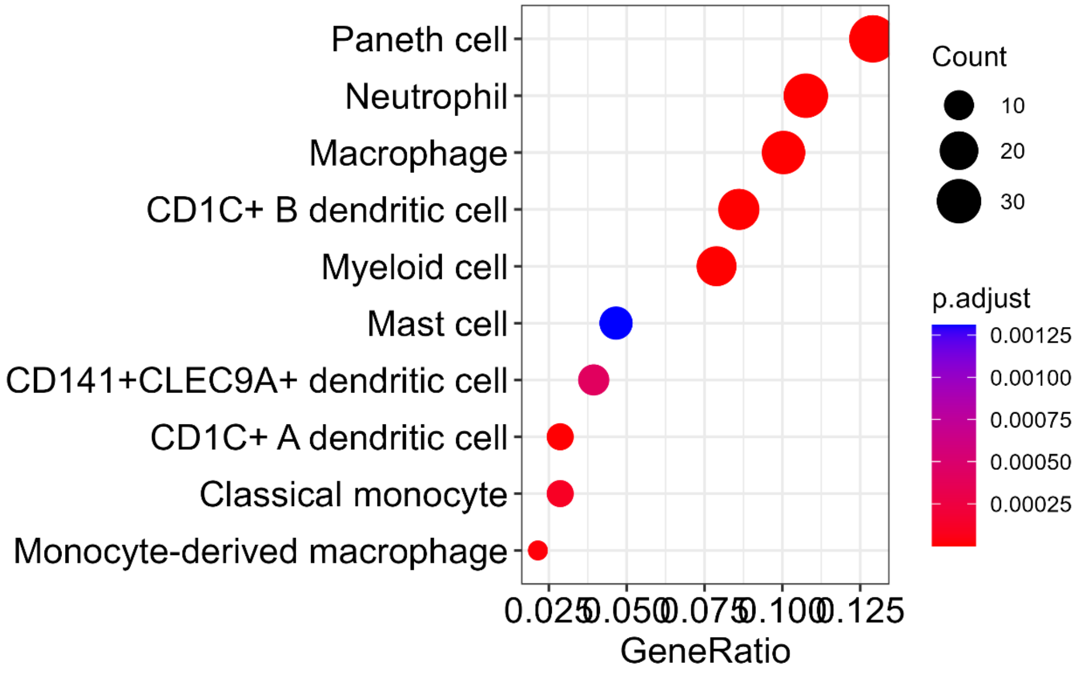


b)

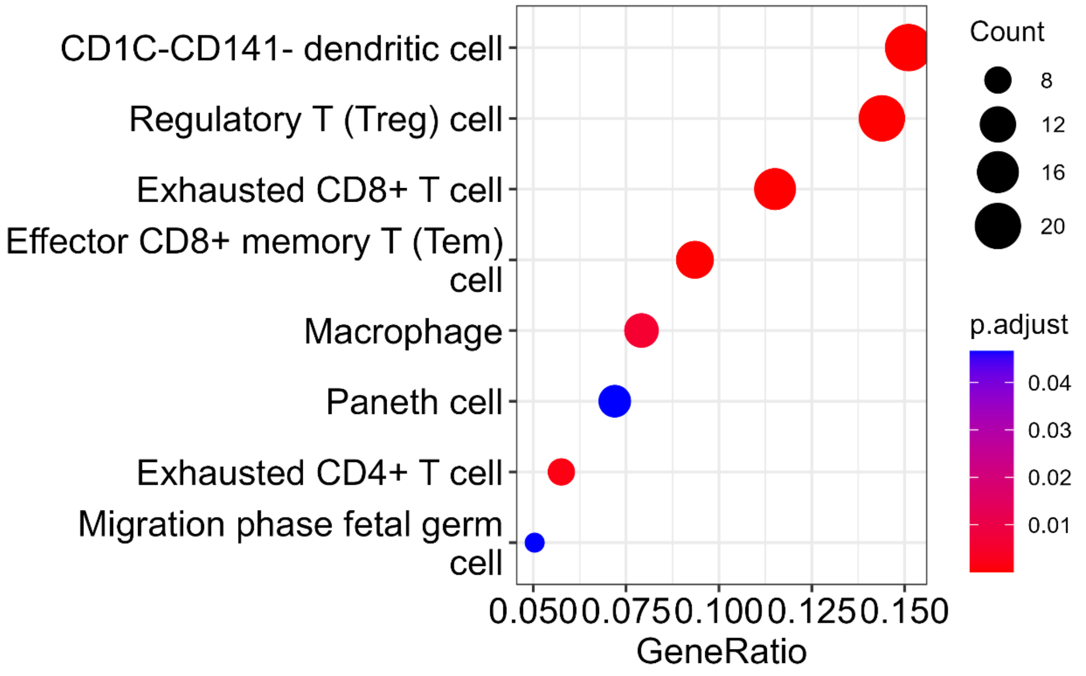

c)

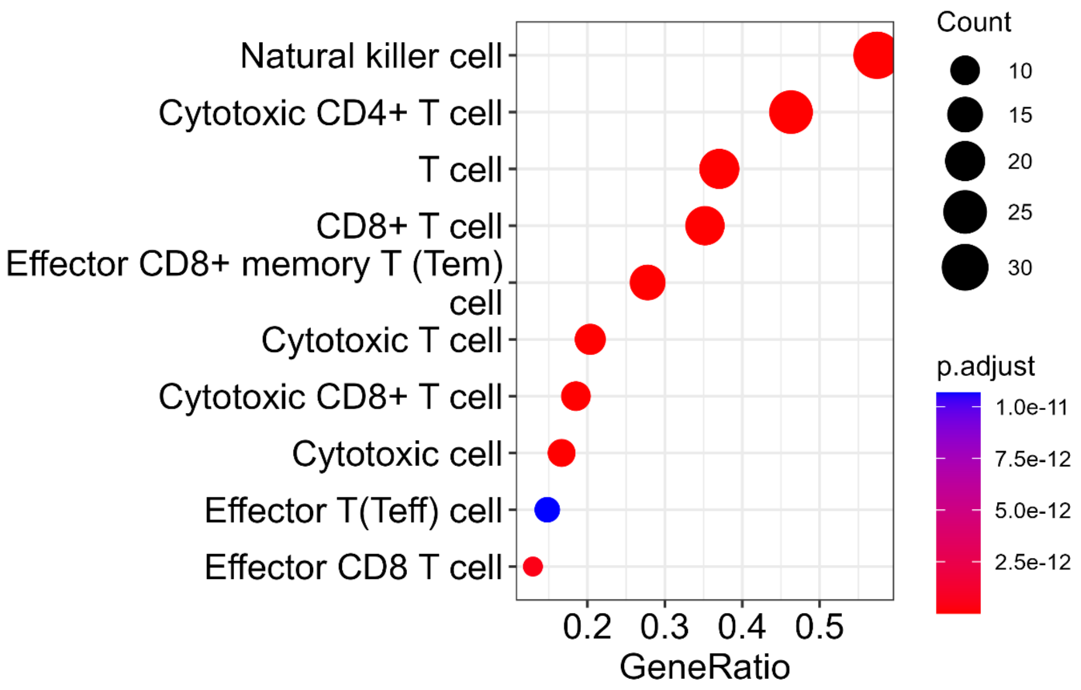
Natural killer cell
Cytotoxic CD4+ T cell
T cell
CD8+ T cell
Effector CD8+ memory T (Tem) cell
Cytotoxic T cell
Cytotoxic CD8+ T cell
Cytotoxic cell
Effector T(Teff) cell
Effector CD8 T cell
0.2
0.3
0.4
0.5
GeneRatio
Count
10
15
20
25
30
p.adjust
1.0e-11
7.5e-12
5.0e-12
2.5e-12

d)

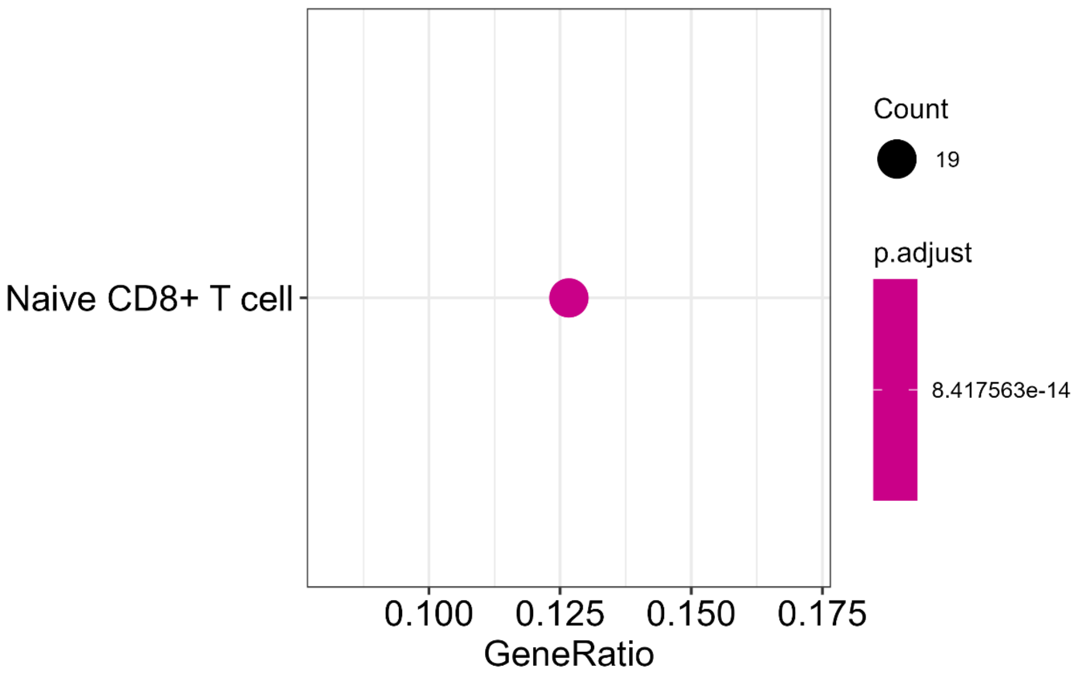
Naive CD8+ T cell
0.100
0.125
0.150
0.175
GeneRatio
Count
19
p.adjust
8.417563e-14

**Fig. S5:** (a) UMAPs visualization of cell clusters from Skin-SHARE data based on scRNA-seq data using ground truth labels as well as labels from MOFA+, WNN, MOJITOO, and TriTan. (b) The clustered heatmap of the correlation matrix, obtained from RNA and ATAC association matrices, showing clusters of gene sets and their potential regulating peak sets. (c) Clustered heatmap of the enrichment matrix from PEGS analysis, x-axis are the gene sets, and y-axis shows peak-sets (expanded to +/-2kb) with the same order of correlation matrix.

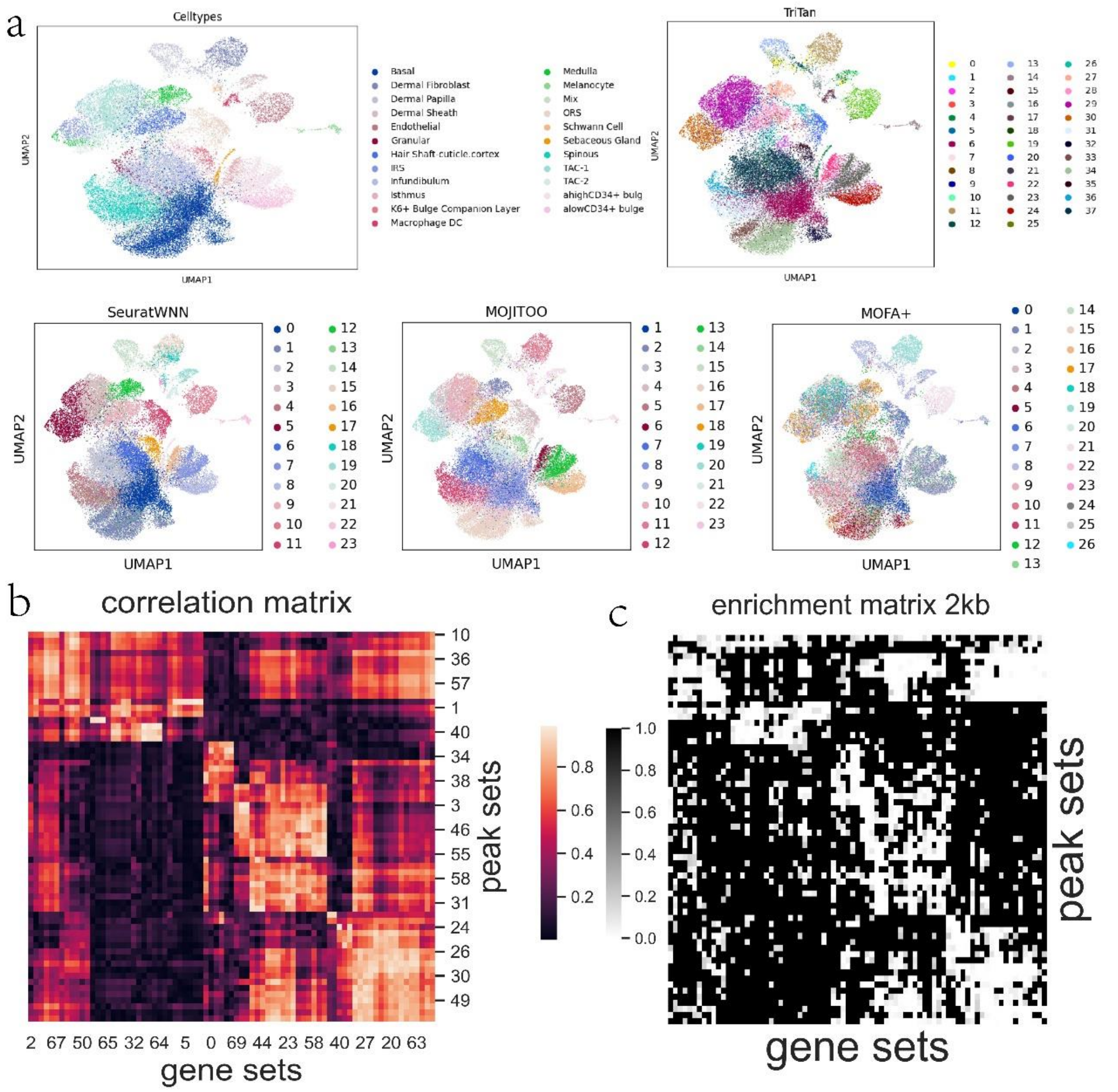

**Table S1:** Gene list of PAX5 regulon in Pro-B cells.

ADAM19 AIM2 ANGPTL1 ANKRD33B ARID5B ATP2A3 BACE2 BANK1 BICD1
BIRC3 BLK BTNL9 C11orf80 CD19 CD1C CD24 CD37 CD79A CD82 CDK14
CELSR1 CHD7 CLEC17A CNR2 COCH COL4A3 COL4A4 CPNE5 CR2 CXXC5
DENND5B DIPK1A EBF1 EML6 FAM30A FCMR FCRL2 FCRL5 FCRLA
FDFT1 FUT8 GALNTL6 GEN1 GNB5 GPM6A HIP1R IFNG-AS1 IGHA1
IGHA2 IGHG1 IGHG2 IGHG3 IGHGP IKZF3 IL7 ISG20 JADE3 LAMC1
LINC01588 LINC01781 MGAT5 MS4A1 NUP88 OSBPL10 P2RX5 PARP1 PARP15
PAWR PDLIM1 PKIG PLEKHG1 PNOC POU2AF1 PPM1K PPP1R16B PTPRG
RABEP2 RALGPS2 RASGRP3 RRAS2 SNX22 SNX25 SOX5 SP140 SSPN
SWAP70 TENT5C TLE1 TLR10 TMEM156 TNFRSF13B TNFRSF13C TRAF5
TSBP1-AS1 UGT8 UST VOPP1 VPREB3 XKR6 ZHX2 ZNF860

**Table S2:** Gene list of PAX5 regulon in Pre-B cells.

SLC2A5 DLGAP3 LINC01362 VAV3-AS1 FCRL1 PLD5 DDAH1 NTNG1 BEND5
ROR1 PIK3C2B FAM177B FCRL3 ENAH GDF7 EHD3 ADD2 PAIP2B
GALNT14 TMEM163 MYO1B NPHP1 CHST10 RGPD2 SLC38A11 LINC01857
LINC01320 LINC02576 MAP2 SCN3A CHL1 LINC01811 KCNH8 KBTBD8
CD200 CD80 BTLA SLC9C1 ZDHHC19 FANCD2OS THRB FAM43A
SERPINI1 TP63 EGOT SLC15A2 LRRC34 INKA1 OSBPL10-AS1 ZDHHC23
CCDC191 LINC01266 GRAMD1C LINC01215 STAP1 PARM1 STPG2 DCLK2
MAPK10 CLNK RASSF6 MMRN1 BEND4 CCSER1 GYPE ANK2 CD180 EPB41L4A
LINC01340 LIX1-AS1 LIX1 SLC23A1 ZNF608 MEF2C-AS1 MEF2C-AS2 COL19A1
MOXD1 TREML2 TXLNB SOBP KHDRBS2 HLA-DQA2 HLA-DQB2 HLA-DOB
HLA-DOA ZNF165 CCR6 PPIL6 TSBP1 AFDN SERPINB9P1 CDCA7L FAM3C
MACC1 CNTNAP2 ABCB4 TSPAN33 RAPGEF5 STEAP1B STAG3 DNAH11 IL6
PYCR3 PEBP4 CLVS1 SLCO5A1 E2F5 TSPYL5 PLPP5 SPATC1 TP53INP1
TPD52 RHOBTB2 ADAM28 ST18 SYBU PAX5 PNPLA7 LCN10 FBXO10
NIPSNAP3B AK8 CD72 GLIDR QSOX2 ADD3-AS1 HSPA12A PAOX KCNIP2
SYT9 SPON1-AS1 ARHGAP42 SLC35F2 DSCAML1 FAM111B LARGE2 FADS3
WEE1 RAB30 NAV2 JAM3 CXCR5 GLYATL1 STK33 SHMT2 TMEM19 SYCP3
GLI1 BCL7A LPAR5 HVCN1 TCTN1 DTX1 BHLHE41 RAD51AP1 LINC02422 SYT1
PCDH9 GAS6-AS1 LINC02324 AKAP6 TCL1A SAV1 CBLN3 MAP3K9 DPF3
IGHD IGHM PRKD1 TCL6 GATM MYO5C TEX9 KNL1 LINC00926
CORO2B SYT17 FA2H PMFBP1 C16orf74 PYCARD-AS1 ATP2A1 DBNDD1 ACSM1
TNFRSF17 GRAPL MYO1C ARHGAP44 EPN2 CCDC144NL-AS1 KLHL14 NETO1
MYO5B RHPN2 SLC6A16 CD22 SYT5 CD70 C19orf18 CCDC106 ZNF418
CNFN FCER2 CACNA1A PPP1R14A ZNF154 MYBPC2 ZNF135 ZBTB32 ACP5
LINC00665 DOCK6 ZNF528-AS1 EYA2 COL9A3 KCNG1 LINC00494 CD40
MACROD2 TPTE ICOSLG SYN3 MGAT3 MICAL3 GUCD1 LARGE1 IGLV10-54
KLF8 XACT CLCN4 PTCHD1-AS WWC3-AS1 FRMPD4

**Table S3:** Parameter values used in Scanpy for selecting highly variable features for three data sets used in the manuscript.

| Dataset | Modality | min_mean | max_mean | min_disp | n_bins |
|---|---|---|---|---|---|
| PBMC-10k | scRNA-seq | 0.0125 | 3.5 | -1 | 20 |
| | scATAC-seq | 0.0002 | 3 | -5 | 20 |
| NeurIPS 2021 | scRNA-seq | 0.0125 | 1.2 | -1 | 1000 |
| | scATAC-seq | 0.0001 | 0.0035 | -2 | 1000 |
| Skin-SHARE | scRNA-seq | 0.0125 | 1 | -1 | 1000 |
| | scATAC-seq | 0.00005 | 0.002 | -2.5 | 1000 |

**Table S4:** Number of SVD components used in TriTan's factorisation for three datasets in the manuscript.

| Dataset | Modality | cell | feature |
|---|---|---|---|
| PBMC-10k | scRNA-seq | 50 | 20 |
| | scATAC-seq | 50 | 20 |
| NeurIPS 2021 | scRNA-seq | 300 | 50 |
| | scATAC-seq | 300 | 50 |
| Skin-SHARE | scRNA-seq | 300 | 50 |
| | scATAC-seq | 300 | 50 |
